# Functional genomic map of local adaptation in sorghum to guide allele mining

**DOI:** 10.64898/2026.05.17.725773

**Authors:** Yuxing Xu, Aayudh Das, Clara Cruet-Burgos, Geoffrey P. Morris, Jesse R. Lasky

**Affiliations:** Department of Biology, Pennsylvania State University, University Park, PA 16802; Department of Soil and Crop Sciences, Colorado State University, Ft Collins, CO 80521; PAC Herbarium Pennsylvania State University, University Park, PA 16802

## Abstract

Genomic data from genebanks could be exploited to find alleles adapted to target environments for resilience breeding, but it can be difficult to prioritize among the thousands of accessions and millions of genomic variants. There are competing hypotheses for the molecular basis and architecture of local adaptations: e.g. whether *cis-*regulatory versus amino acid changing variants are more important; or whether small-effect, low pleiotropy versus large-effect, high pleiotropy variants are more important. Here, we compare a range of variant types and genomic contexts thought to influence effect size, pleiotropy, and selection for their role in local adaptation in 443 whole genome resequenced African sorghum landraces. We used genotype-environment associations (GEAs) as evidence of local adaptation. We found that GEA were particularly enriched in the vicinity of genes and depleted elsewhere. However, enrichment was strongest in likely *cis*-regulatory contexts: accessible chromatin, unmethylated regions, and in transposable elements close to genes. Near genes, there were clear peaks in GEAs at the transcription start site, where mutations are demonstrated to have the largest expression effects. Additionally, GEAs in accessible chromatin and unmethylated regions were better predictors of genetic variation in response to experimental drought than comparable loci. Having tested hypotheses about the variants underlying local adaptation, we can now use this knowledge of the importance of *cis*-regulatory variation in the search for new environmentally-adaptive alleles for plant improvement.

## Introduction

Global crop genebanks house vast collections of diverse germplasm thanks to the foresight of agricultural researchers who collected and stored germplasm (1, 2). At Vavilov’s genebank during the siege of Leningrad (from 1941-1944) and resulting famine, staff took great efforts to protect their collection, even at the cost of their own lives (3). Why go through such effort? Genebanks have long been promised as genetic warehouses for breeding (4). Now, an urgent challenge is to breed crops for future environments. Fortunately, traditional local varieties from around the world (“landraces”) of crops are typically adapted to diverse local environments (5, 6). Local adaptation is defined as a genotype-by-environment interaction for fitness where native genotypes outperform nonlocal ones (7). However, a challenge in leveraging genebanks is the overwhelming size of the collections, numbering in the tens of thousands for major crops. Genomic data from these accessions could help in mining useful alleles (8). If researchers following up on forward genetic screens or looking for adaptive variants in reverse genetics candidates knew which types of mutations and genes are most important for local adaptation, then the search could be more efficient (9). What genomics information could be used to move adaptive alleles from seed bank to seed bag, i.e. into farmers hands and fields?

Eukaryotic genomes are littered with diverse features (some only recently discovered (10)) that can affect phenotypes by altering gene regulation, translation, protein sequence, and folding. Previous debates on the molecular basis of adaptation often centered on the role of *cis-* regulatory versus amino acid variants (11, 12). The hypothesis that *cis-*regulatory variants are most important in adaptation was founded on the assumption (still rarely tested) that these mutations were likely to be less pleiotropic and of smaller effect size than protein altering mutations, which have a greater chance of disrupting protein function across all contexts where a gene is expressed, leading to negative pleiotropy (Fig. 1A) (13). Theory on local adaptation has often focused on effect size and pleiotropy, e.g. finding that gene flow between environments will favor a genetic architecture concentrated in large effect variants (14). More evidence is required to determine the empirical roles in local adaptation of variants that are protein coding, *cis*-regulatory, and found across the diverse features of genomes.

**Figure 1.**
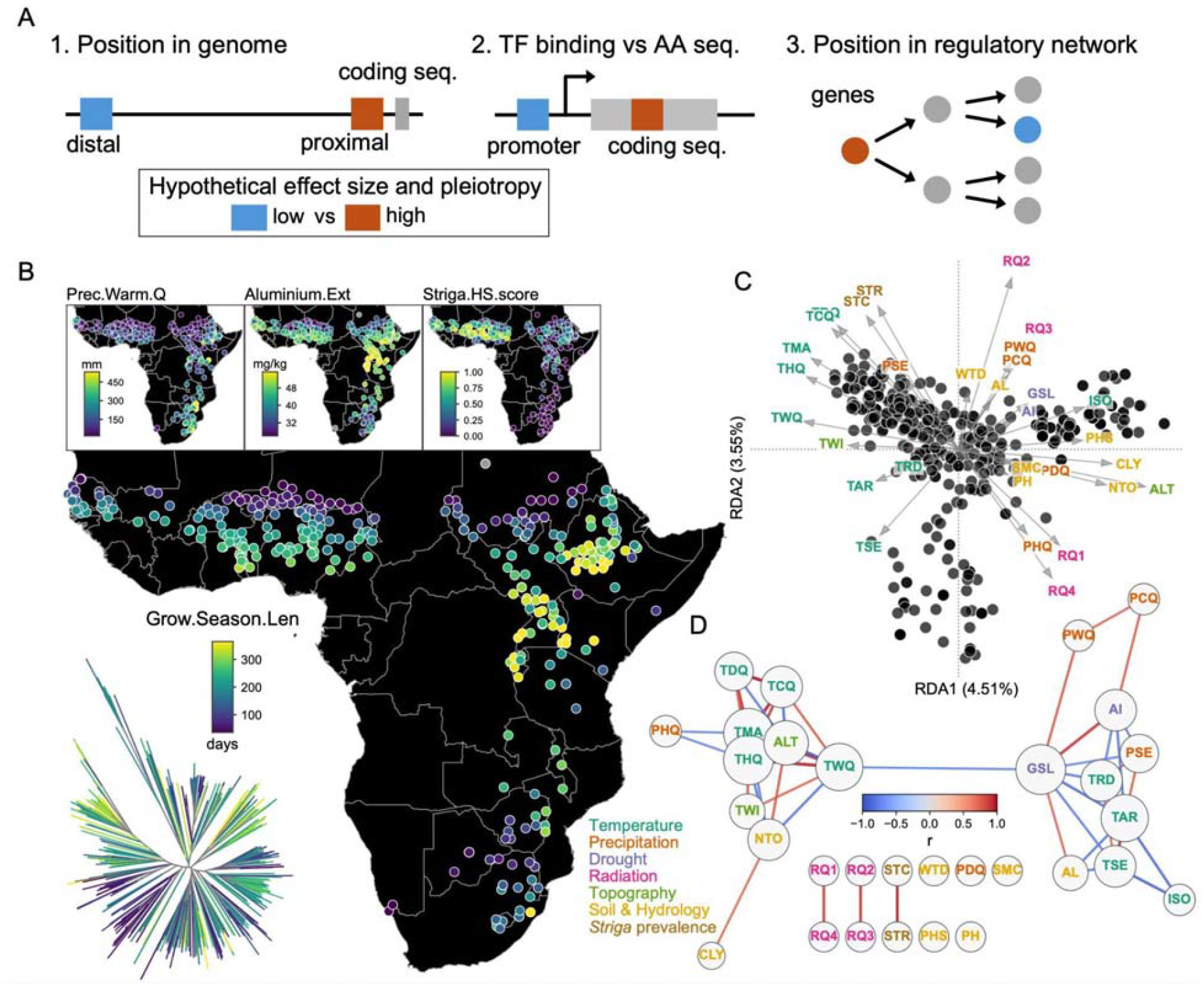
Competing hypotheses for the genetic basis of local adaptation (A) can be tested in African sorghum landraces from diverse environments (B-D). (A) Example competing hypotheses about effect size and pleiotropy of adaptation: (1) genomic compartment (genic vs transposable element/repetitive sequence), (2) functional element within or near genes (regulatory vs coding; e.g., promoter, coding sequence, intron), (3) regulatory-network position (upstream vs downstream). Colors denote low versus high expected effect size and pleiotropy. (B) Geographic distribution of African sorghum landraces and four representative environmental variables used in genotype–environment association (GEA) analyses: precipitation of the warmest quarter (PHQ), extractable aluminium (AL), Striga habitat suitability score (STR), and growing season length (GSL, a proxy for terminal drought). Points denote landraces and are colored by environment. The neighbor-joining tree (lower left) shows landraces, with tip colors indicating growing season length. (C) Distribution of sorghum landraces along the first two axes of redundancy analysis (RDA). Arrows indicate the direction and relative contribution of environmental variables. (D) Correlation network of environmental variables. Nodes represent environmental variables, with node size reflecting degree within the network, and edges indicating correlations between variables, with edge color reflecting correlation direction and magnitude (|r|>0.7 shown). In (C) and (D), environmental variables are colored by category.

One challenge for studying the molecular basis of adaptation is that while the genetic codes for amino acid sequences of proteins are well understood, gene regulation is a more complex (arising from multiple interacting forces) and noisy (e.g. the tolerance of transcription factors for variable binding sequences) process (15). However, new functional genomics assays are allowing identification of *cis-*regulatory sequences (16–19). Even the effects of amino acid substitutions on protein function are challenging to predict (20, 21), but recent advances permit computational predictions based on e.g. sequence conservation (22). At another biological level, the challenge of characterizing the ecology of selection can be approached with genotype-environment associations (GEAs) (9, 23). GEA genome scans identify loci where allele frequency is strongly correlated with environment, which is evidence of local adaptation. GEAs have been frequently employed (24), including for crop landraces (25–28). However, GEAs are subject to false positives and negatives, and should be complemented with phenotypic data (9, 29, 30). Our goal is to determine what types of variants are most important in local adaptation, and use this knowledge to inform breeding for crop resilience.

Sorghum is a globally cultivated cereal important to smallholders in tropical regions, and shows great genetic variation in phenotypes among landraces from different environments (27, 31–33). We have previously shown in sorghum that loci identified from GEA can predict adaptation to drought and soil aluminum toxicity, and that known quantitative trait loci (QTL) for adaptation to climate, photoperiod, and *Striga* parasitism show strong GEAs (27, 34). Here, we use environmental genome-wide association studies to identify locally adapted loci in sorghum landraces, focusing on drought, aluminum, and *Striga* adaptation. We tested for enrichments of GEAs in a variety of functional contexts. We show evidence that *cis-*regulatory elements are important in local adaptation, with enrichments peaking at the beginning and end of transcription, accessible chromatin, and unmethylated regions. These findings can help prioritize candidate alleles from sequenced genebanks for experiments and breeding.

## Results

To identify locally adapted variants, we used a high-depth whole-genome resequencing of 443 georeferenced African sorghum landraces sampled across diverse agroecological regions (mean depth 43.4×, Dataset S1) (32). This panel includes >15 recognized botanical types or intermediates (SI Appendix, Fig. S1) and captures major environmental gradients across African sorghum-growing regions (Fig. 1B). The initial variant discovery identified 27,945,979 single-nucleotide polymorphisms (SNPs) and 1,861,758 insertions/deletions (InDels) (32). For subsequent association analyses, we retained variants with minor allele frequency (MAF) ≥ 0.05 and heterozygous genotypes present in no more than 5% of accessions (avoiding likely mapping errors), yielding 5,930,896 high-quality SNPs and 521,682 InDels. We compiled 31 environmental variables for each landrace, spanning climate means, seasonality, soil properties, solar radiation, and the prevalence of the parasitic plant *Striga hermonthica* (Fig. 1C, SI Appendix, Table S1).

To assess whether environmental gradients explain genome-wide variation in African landraces, we performed redundancy analysis (RDA) using 31 environmental variables and 987,993 unlinked SNPs. Environment explained 18.7% of genomic variation, with the leading axes mainly reflecting temperature/elevation and precipitation/radiation gradients (Fig. 1C). To characterize GEA across the full environmental dataset, we performed environmental genome-wide association study (envGWAS) for each variable using linear mixed models and found that signal profiles were highly correlated among related variables, mirroring the environmental correlation structure (Fig. 1D; Fig. 2F). To reduce redundancy while retaining literature-supported stress gradients, we focused analyses below on four variables (SI Appendix, Figs. S2-S5): growing season length and precipitation of the warmest quarter as complementary drought measures (27), soil extractable aluminium as a proxy for aluminium toxicity (27), and a *Striga* habitat suitability score as a proxy for parasite pressure (34).

**Figure 2.**
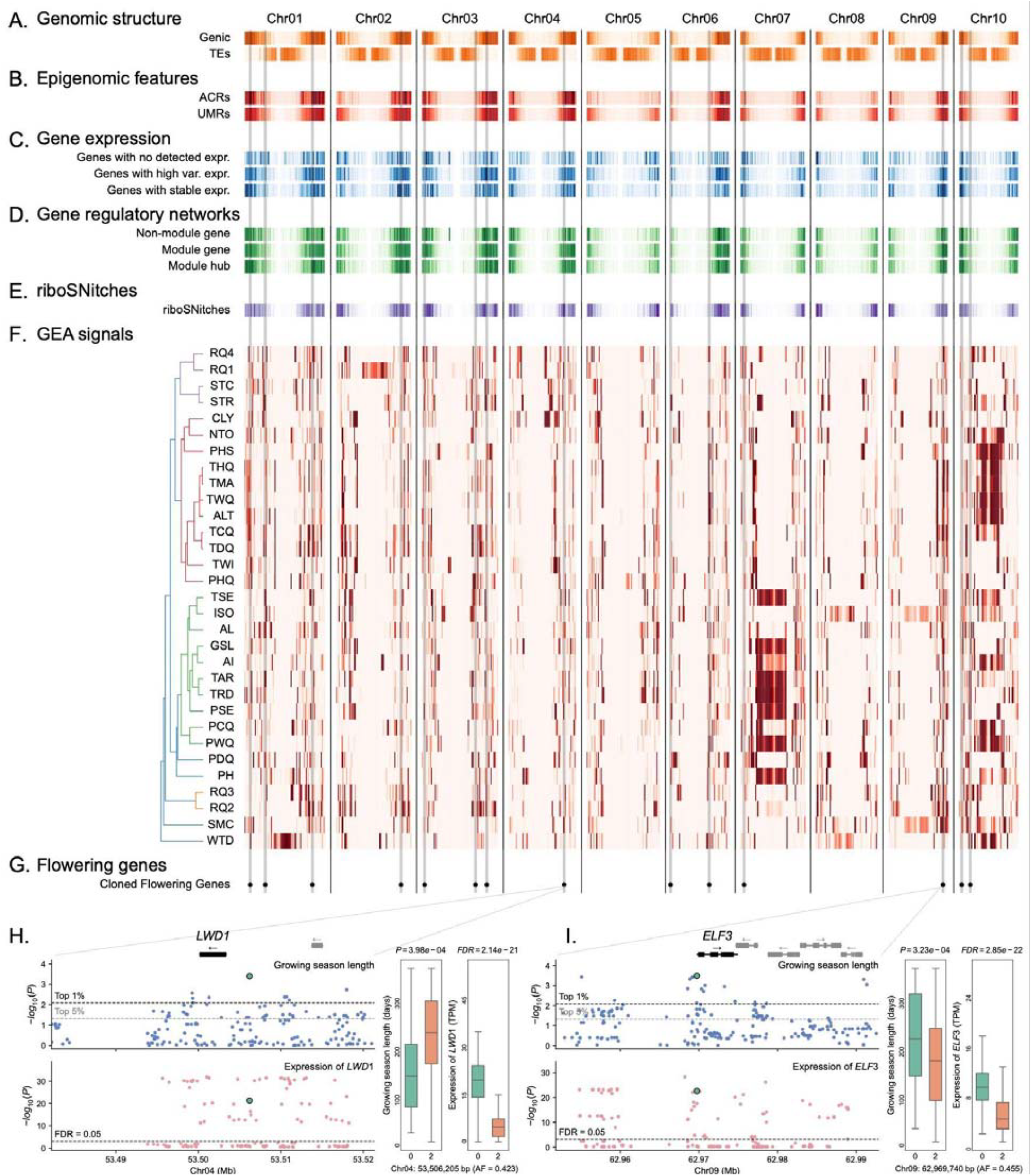
The diverse landscape of genomic features and genotype-environmental associations (GEAs). The ten sorghum chromosomes (Chr01–Chr10) were divided into equal-length bins. All tracks were z-scored within track; darker cells indicate higher-than-mean values, lighter cells indicate lower-than-mean values. (A) Genomic features: gene density, transposable-element density. (B) Epigenomic features: accessible chromatin regions (ACRs) and unmethylated regions (UMRs). (C) Gene-expression categories: genes with no detected expression, genes with high expression variance, and genes with stable expression. (D) Gene regulatory network categories: non-module genes, module genes, and module hubs. (E) Predicted riboSNitches. (F) envGWAS: Each row corresponds to one environmental variable. The dendrogram clusters variables by similarity of their per-sample values across the resequenced landrace panel. The heat map shows enrichment of the top 5% significant SNPs across bins, calculated with a hypergeometric test using callable SNPs per bin as background, expressed as −log10(P), and z-scored within row. Darker cells indicate relative enrichment for that variable. Flowering-time genes are indicated along the x-axis; gray vertical bands span panels. (G) Genomic positions of cloned flowering-time genes. Gray vertical lines indicate the corresponding chromosome positions and extend through the upper genome-wide tracks. (H–I) Locus zooms for two flowering-time genes. Gene models and genomic coordinates are shown at top. The upper (blue) plots display SNP associations between genotype and the growing season length (GSL) environmental variable; light and dark dashed lines indicate the genome-wide envGWAS top 5% and top 1% thresholds, respectively. The lower (red) Manhattan plots show associations between SNP genotype and the expression of the focal gene (putative cis-eQTLs); the dashed line denotes FDR=0.05.

### Genotype environment associations are enriched in flowering time genes

Appropriate flowering time for local environments and farmer preferences is a critical trait for crops (35, 36), and local adaptation often involves flowering time in wild species (26, 37–39). To assess whether GEAs are capturing true local adaptations, we compared drought associated loci with those affecting flowering time. We defined a sorghum flowering-time gene set by identifying orthologs of Arabidopsis flowering-time genes in the FLOR-ID database (40), yielding 312 genes (Dataset S4). We tested whether flowering time genes were enriched in GEAs (Methods) and found these genes most enriched in GEAs for growing season length (*P*=0.009; SI Appendix, Fig. S6 and Table S3), a variable reflecting seasonal drought, and where rapid flowering in short growing seasons allows drought escape (41). In contrast, no enrichment was detected for precipitation of the warmest quarter (*P*=0.25, SI Appendix, Fig. S6) or for variables unrelated to drought (Aluminum and *Striga*, SI Appendix, Fig. S6).

We tested if the canonical flowering time genes had likely *cis-*regulatory variants playing a role in local adaptation because some hypotheses posit that adaptation tends to involve *cis-* regulatory variants (12, 13) (SI Appendix, Dataset S4). We found several genes where *cis-*eQTL had allele frequency clines along climate gradients, including orthologs of core circadian and photoperiod regulators *LWD1/2* (LIGHT REGULATED WD1/2, Fig. 2H) (42), *ELF3* (EARLY FLOWERING 3, Fig. 2I) (43), *TEM1* (TEMPRANILLO 1) (44), and *CHE* (CCA1 HIKING EXPEDITION) (45). *ELF3* for example has been identified as a QTL for photoperiod response in sorghum, a critical trait for local adaptation (46).

### Local adaptation signals are enriched in and around expressed genes, including gene-proximal transposable elements

Crop geneticists sifting through putative QTL from forward genetics screens of genebank accessions may intuitively pursue variants that are in or near genes as opposed to those distant from genes. This strategy is based on a hypothesis that most adaptations involve changes to amino acid sequences or nearby linked *cis-*regulatory elements, suggesting stronger adaptation signals in genic sequences (47). We found that genic SNPs were enriched in associations with all four environmental variables (precipitation of the warmest quarter, *P*=0.014; growing season length, *P*=0.020; extractable aluminium, *P*=0.006; *Striga* HS score, *P*=0.017; *z*=2.0–2.7; Fig. 3A; SI Appendix, Table S2).

**Figure 3.**
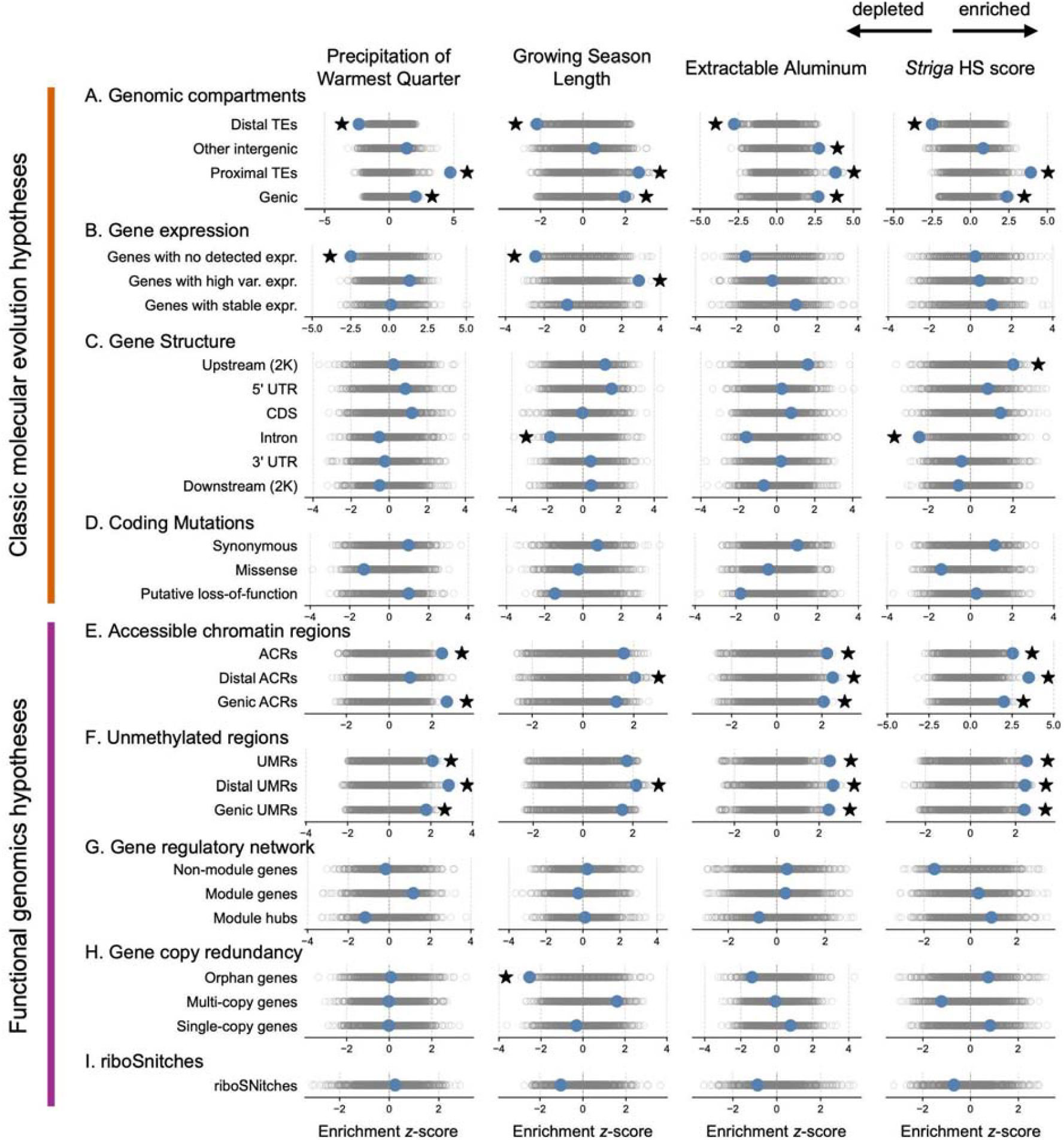
Enrichments of GEAs across categories of functional variation. Hypotheses for the contexts and characteristics of locally adaptive variants are divided into “Classic molecular evolution hypotheses” (A-D) and “Functional genomics hypotheses” (E-I). Enrichments were tested on the top 5% of GEAs. Observed values (blue) are calculated as a z-score using 1,000 null permutations (gray). Significant enrichments are indicated by stars (two-sided empirical P<0.05).

Some annotated genes may be pseudogenes or very rarely expressed, which we hypothesized would tend to not play a role in adaptation. Consistent with this hypothesis, we found non-expressed genes were generally depleted in environmental associations, significantly in the case of precipitation of the warmest quarter (*P*=0.003; *z*=-2.5) and growing season length (*P*=0.001; *z*=-2.5; Fig. 3B; SI Appendix, Table S3).

There are prominent examples in the literature of transposable elements (TEs) fueling adaptation, often by gain or loss of *cis*-regulatory elements, such as a TE insertion in the 5’UTR of wheat *VRN1* that disrupts its vernalization-dependent regulation, but not the *VRN1* coding sequence, permitting a spring annual habit (48–50). Concordantly, we found the TEs proximal to genes (<5 kb from a gene) were highly enriched in GEAs (precipitation of the warmest quarter, *P*=0.001; growing season length, *P*=0.012; extractable aluminium, *P*=0.006; *Striga* HS score, *P*=0.001; *z*=2.6–4.7; Fig. 3A). These strong signals of local adaptation at proximal TEs could indicate particular environments select for *cis-*regulatory mutations arising subsequent to the TE insertion or that the insertion itself is polymorphic (though present in the reference genome) and its insertion is locally adaptive with the SNPs we tested tagging the insertion.

Most TEs, however, are not proximal to genes and less likely therefore to be regulatory. Most are found in pericentromeric regions (51) (SI Appendix, Fig. S7) where low recombination limits effective population size and may constrain adaptation ((52), but see (53, 54)). Accordingly, we found distal TEs (i.e. >5 kb from annotated genes) were depleted in association with all four environmental variables (precipitation of the warmest quarter, *P*=0.004; growing season length, *P*=0.007; extractable aluminium, *P*=0.002; *Striga* HS score, *P*=0.001; *z*=−2.2 to −2.8; Fig. 3A). Overall these results support the hypothesis of adaptation primarily in and near genes, suggesting coding or *cis-*regulatory variants may be most useful for crop environmental adaptation.

### No significant signal of local adaptation comparing missense, nonsense, and synonymous variants

A classic hypothesis is that missense (non-synonymous) mutations more likely have phenotypic effects and hence are subject to stronger positive or purifying selection than neighboring synonymous mutations (55). This hypothesis has been tested multiple times in the context of GEA and local adaptation in crop landraces (27, 56) wild relatives (57) and other plants (38, 58, 59), with mixed results, though few previous results come from high coverage resequencing data. Here, we found no significant patterns across all four variables; synonymous SNPs were only slightly enriched and missense SNPs only slightly depleted in GEAs (synonymous: *z*=0.8−1.1, *P*=0.116–0.211; missense: *z*=−1.4 to −0.2, *P*=0.600–0.912; Fig. 3D).

In addition to missense and synonymous SNPs, other coding mutations could disrupt the protein as a whole (e.g. frameshifts), mutations classically thought to be deleterious (60). However, there are prominent examples of adaptation by gene loss of function, including local adaptation in crop landraces, like the loss of function in sorghum *LGS1* in regions prone to *Striga* infestation (34, 61–63). Additionally, these mutations may be more likely to have a phenotype than the average SNP across the genome (most of which probably have no phenotypic effect), perhaps suggesting potential for adaptation. Here we found that these putative loss-of-function variants had no clear enrichment in GEAs for all four environmental variables (*z*=−1.8 to 1.0, *P*=0.152–0.969; Fig. 3D). Thus while particular coding variants with phenotypic effects (e.g. missense and nonsense SNPs) may be important to local adaptation, globally consistent purifying and positive selection (limiting GEAs) may be more important in determining spatial variation in coding sequences.

### Sites with likely cis-regulatory epigenomic features are highly enriched in signals of local adaptation

*Cis-*regulatory evolution underlies some prominent adaptive traits in crops but its role in local adaptation genome-wide has not been explored in depth. The original hypotheses of the importance of *cis-*regulatory adaptation (12, 13) predate the high throughput assays that now permit identification of likely regulatory sequences across the genome. Accessible chromatin is a good marker of where transcription factors may bind, and high throughput assays can find accessible chromatin regions (ACRs). We used published sorghum leaf ACRs (17) and found that these were significantly enriched in GEAs for three of the four environmental variables (precipitation of warmest quarter: *z*=2.5, *P*=0.001; extractable aluminium: *z*=2.2, *P*=0.003; *Striga* HS score: *z*=2.5, *P*=0.003; Fig. 3E). These enrichments were consistent when considering only gene-proximal ACRs (≤2 kb from genes; precipitation of warmest quarter: *z*=2.7, *P*=0.001; extractable aluminium: *z*=2.0, *P*=0.007; *Striga* HS score: *z*=2.0, *P*=0.015) or only gene-distal ACRs (>2 kb from genes; extractable aluminium: *z*=2.5, *P*=0.004; *Striga* HS score: *z*=3.5, *P*=0.002), suggesting that both promoter-proximal and more distal *cis*-regulatory regions are hotspots of local adaptation, though distal ACRs may include enhancer-like elements as well as unannotated or extended 5′ promoter regions (64).

Chromatin accessibility can change across cell types and environments, and a single ACR dataset may miss many regulatory regions. By contrast, un-methylated regions (UMRs) are consistent across cell type and environment and good indicators of sites that are regulatory in at least some contexts (18). We found that published sorghum UMRs (18) were strongly enriched in evidence for local adaptation and in some cases beyond that found with ACRs (precipitation of warmest quarter: *z*=2.1, *P*=0.009; extractable aluminium: *z*=2.5, *P*=0.004; *Striga* HS score: *z*=2.5, *P*=0.005; Fig. 3F). Again we stratified by UMRs proximal and distal to genes, and found these both were significantly enriched in GEAs (proximal UMRs: precipitation of warmest quarter: *z*=1.8, *P*=0.018; extractable aluminium: *z*=2.5, *P*=0.004; *Striga* HS score: *z*=2.4, *P*=0.006; distal UMRs: *z*=2.1–2.9, *P*=0.003–0.020; Fig. 3F). Thus, UMRs may be good markers of locally adapted *cis-*regulatory loci in sorghum landraces.

Local adaptation in crops may be highly polygenic and best modeled with genomic relatedness matrices (GRMs) (27, 65, 66). Because of the strong enrichments of GEAs within UMRs, ACRs, and proximal TEs, we also tested how much variance in environmental variables could be explained by GRMs built using SNPs in each genomic context. We compared the observed variance explained with that explained using a set of matched loci (Fig. 4B) and found that ACRs, UMRs, and proximal TEs explained more variance than expected for most context–environment combinations (9/12 significant; exceptions: ACRs for precipitation of the warmest quarter and growing season length, and proximal TEs for growing season length), indicating that environment-associated genetic variation was concentrated in these genomic contexts. The strongest pattern was observed for UMRs, where sites explained significantly more variance than matched random site sets for all four environmental variables (*P*≤0.001, *z*=9.4–18.3; SI Appendix, Table S4). These results suggest that GRMs built with UMRs (and to a lesser degree ACRs, proximal TEs) could be useful for genomic selection/prediction approaches to identify useful genotypes in genebanks.

**Figure 4.**
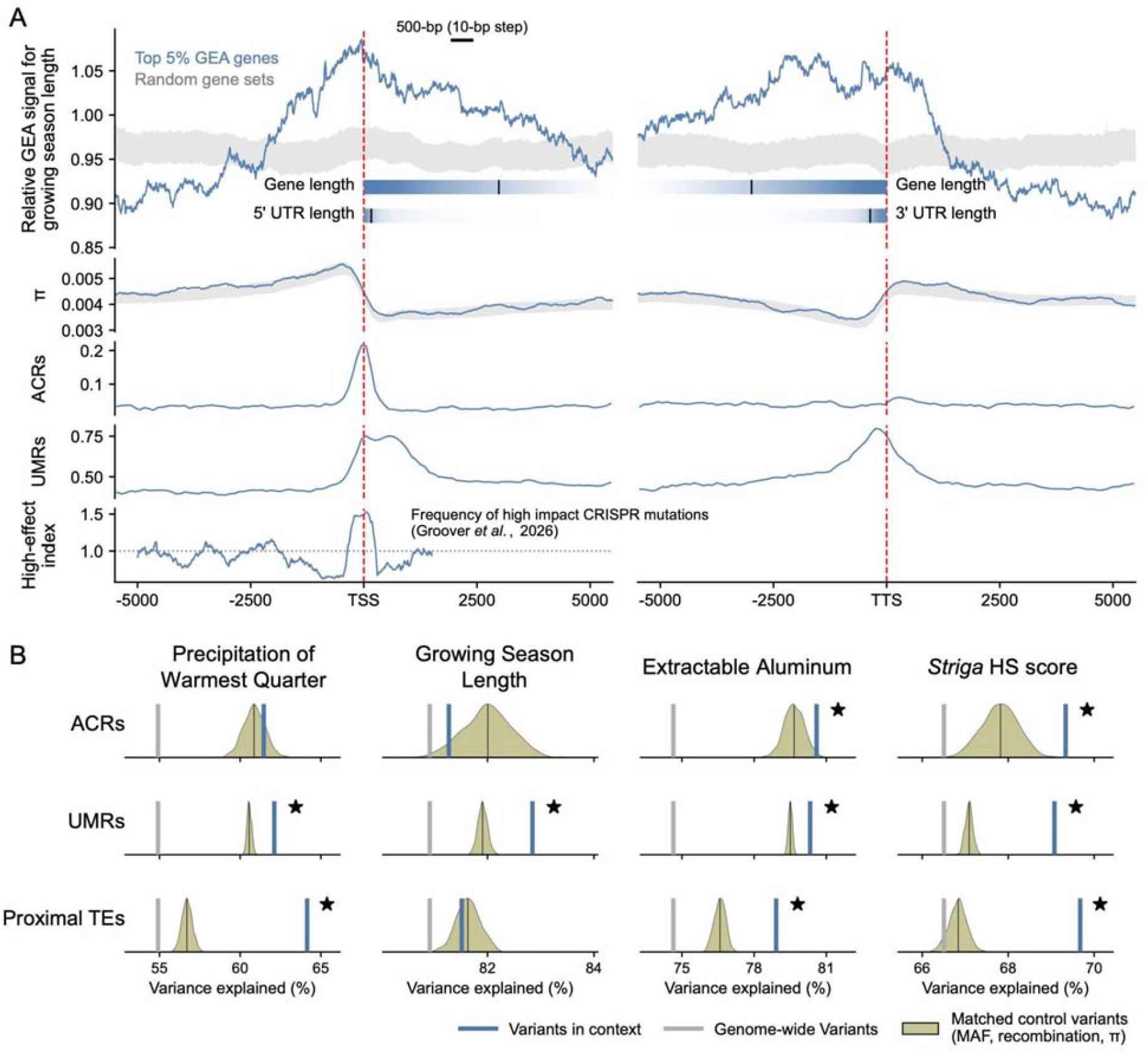
Cis-regulatory regions are enriched for GEAs and explain more environmental variance than the genomic background. **(A)** Spatial distribution of envGWAS signals around genes associated with growing season length. The panel shows signal profiles around both ends of the top 5% of genes most strongly associated with growing season length. The x-axis indicates distance (bp) from the transcription start site (TSS; left) or transcription termination site (TTS; right). The bars labeled gene length and 5′/3′ UTR length show the distribution of gene and UTR lengths across genes, where darker shading indicates a higher proportion of genes extending to that position and black vertical ticks indicate the median length. The dark-blue curve shows the relative enrichment of envGWAS signals, calculated as the ratio of the first quartile (Q1) of −log10(P) values within sliding windows (500 bp window, 10 bp step) to the corresponding Q1 across the entire ±10 kb region around the TSS or TTS; the grey band denotes the 95% empirical range from 1,000 genome-wide random gene sets of equal size. The ACR and UMR tracks represent the proportion of bases overlapping each feature, and π shows nucleotide diversity in the same windows. The bottom track shows a high-effect index, measuring the enrichment of high-effect CRISPR-accessible promoter mutations in each window from Groover et al. (2026) (dotted line, index=1: no enrichment). **(B)** Variance explained by sites within genomic contexts. Columns correspond to environmental variables and rows correspond to genomic contexts (ACRs, UMRs, and proximal TEs). Genomic relationship matrices (GRMs) were constructed using genotypes at sites within each context, and the proportion of variance in each environmental variable explained by each GRM was estimated. Blue vertical lines indicate the variance explained by sites within each genomic context, grey vertical lines indicate the variance explained by genome-wide sites, and beige distributions show null expectations from GRMs constructed using randomly sampled sites matched to the context sites by minor allele frequency, recombination rate, and nucleotide diversity (π) across 1,000 replicates.

### Local adaptation signals are strongest at transcription start sites (TSSs)

While the above analyses show genic and putative *cis-*regulatory loci are enriched for GEAs, it is still not clear to what degree selection directly acting on one of the classes of loci may end up driving the statistical signals at linked loci. We first tested separately the categories of sequence near genes: upstream (2 kb), 5′ UTR, CDS, intron, 3′ UTR, and downstream (2 kb). However, possibly due to the linkage between the proximal loci, we found no clear differences among most categories (Fig. 3C), except for the intronic sequence, which showed significant depletion of GEA signals in two environmental variables (growing season length: *z*=−1.8, *P*=0.020; *Striga* HS score: *z*=−2.4, *P*=0.004; Fig. 3C).

To better estimate where adaptation is strongest, we examined the fine-scale distribution of GEAs around genes. Specifically, we aligned the top 5% most significant genes from the gene-level analysis by their transcription start sites (TSSs) and transcription termination sites (TTSs), and quantified local relative signal intensity across the ±10 kb regions surrounding each feature using sliding-window analyses (SI Appendix). We found peaks in GEAs at the TSS, with a lower and broader plateau through genic regions and ∼1 kb before the TSS and after the TTS (Fig. 4A). This pattern was repeated for all four environmental variables (SI Appendix, Figs. S9-S11), suggesting that mutations at and near the TSS have phenotypic effects causing local adaptation. When we aggregated the signal of UMRs across genes, we also found that these peak in frequency just downstream and upstream of the TSS and TTS, respectively, partially aligning with the GEA peaks (Fig. 4A). The portion of GEA enrichment in the gene is clearly distinct from the UMR peaks, however, suggesting a signal of important mutations affecting amino acid sequences and transcript splicing (i.e. mutations in genic sequences). Notably, the peak in GEA enrichment begins its sharpest increase ∼500 bp upstream of the TSS, where UMRs start to increase, and where a recent massively parallel reporter assay showed the greatest *cis*-regulatory effects of small CRISPR mutations in three sorghum photosynthesis genes (67).

Because environmental associations could be spurious, and perhaps biased toward high diversity sites, we evaluated variation in nucleotide diversity (π) in the same region. We found quite distinct patterns of π, compared to GEA enrichment (Fig. 4A), indicating nucleotide diversity alone is not driving the signal we identified.

### No evidence of local adaptation particular to genes at different positions in coexpression networks

Genes in different positions in molecular interaction (e.g. gene regulatory) networks may have different levels of pleiotropy thus have different distributions of mutational fitness effects and patterns of evolution (68–70). For example, drought-imposed selection on rice acts more strongly on peripheral genes (71) while in other plants genes involved in local adaptation can occupy central (72, 73) or peripheral positions in networks (74). We built gene coexpression networks using 1,923 sorghum RNA-seq datasets spanning diverse tissues, developmental stages, and experimental conditions, and tested for GEA enrichment among module, hub, or non-module genes. Across all environmental variables, local adaptation signals were not significantly enriched or depleted among these categories (Fig. 3G; SI Appendix, Table S3), indicating that local adaptation is not preferentially localized to specific positions within the network.

### Variants that alter secondary structure of mRNA not enriched in evidence of local adaptation

RiboSNitches are a relatively recently discovered class of regulatory variants (10) where coding sequence changes alter the folding and secondary structure of mRNA; this altered structure may then affect aspects of mRNA behavior like splicing and translation (75, 76). Because structure is also affected by environment, riboSNitches have been hypothesized to play a role in local adaptation (77). To identify putative riboSNitches across the sorghum genome, we compared the predicted structural ensembles of reference and alternative sequences for natural variants located within protein-coding genes, including introns and UTRs, and classified variants with a correlation coefficient<0.8 as riboSNitches (77). Among 2,517,753 such variants, 696,192 were predicted to be riboSNitches, accounting for 27.65% of all tested sites. This proportion is remarkably similar to that reported in Arabidopsis (77), where 27.11% of transcript-associated SNPs were predicted riboSNitches. Overall, predicted riboSNitches did not show significant enrichment in GEA (all *P*>0.05; Fig. 3I). Thus, while many riboSNitches might be locally adaptive (77), the picture in aggregate is more complex. Additionally, limitations to computational prediction of riboSNitches could limit our ability to discern aggregate signals of selection.

### Variants and genes exhibiting distinct evolutionary dynamics show mixed evidence of local adaptation

Above we primarily focused on testing hypotheses about the roles of protein coding and regulatory variants in local adaptation. These hypotheses were not exhaustive, other hypotheses predict how the evolutionary dynamics of genes and individual variants reflect their effect sizes and pleiotropy. For example, the rate at which genes evolve over deep time may be determined by their function and the effect size of their mutations (78–80). Low conservation sites may be less constrained by negative pleiotropy and may rapidly evolve new locally-adapted functions (81). Alternatively, local adaptation may involve conserved sites because of a greater likelihood their mutations have phenotypic effects (82–84), and some conserved genes may be under spatially varying selection in different species (73). With a few exceptions, however, genes differing in conservation/lability did not show distinct patterns of local adaptation (SI Appendix, Fig. S8). Relatedly, gene families show diverse dynamics, as genes undergo gain, duplication, loss, and functional divergence, and these distinct evolutionary trajectories may influence the fitness effects of mutations in component genes and their role in crop adaptation (85, 86). We found that orphan genes (not in any family) were depleted among genes associated with growing season length (*z*=-2.5, *P*=0.006), suggesting that local adaptation may favor older gene families, but other categories and environmental variables did not show significant enrichments (Fig. 3H; SI Appendix, Table S3).

The frequency and distribution of locally adaptive variants could have tremendous impacts on efforts to discover these variants in genebanks and deploy them in breeding. For example, when different mutations cause adaptation to similar environments in different regions, rarer variants cause local adaptation (87), suggesting large germplasm collections are essential. Alternatively, if local adaptation occurs via the same mutations across a species range, then these variants should be at higher frequency, indicating such variants are already in regional germplasm. We found opposite frequency patterns for different environmental gradients: drought and soil-alumiminum associated variants were significantly uncommon while *Striga-associated* variants were more common and in significantly higher diversity loci (SI Appendix, Fig. S8). These results suggest that differences among environmental gradients (e.g. in geometry) and the biology of adaptive traits may lead to distinct genetic architectures (88, 89), indicating both global and regional germplasm are useful for mining locally-adapted alleles.

### Genotype-environment associations in cis-regulatory contexts better predict genetic variation in drought response (GxE) than control loci

In some cases GEAs can be misleading (90) and thus should be also tested with phenotypic data. We previously showed that GEAs in global sorghum landraces can predict genetic variation in response to managed drought (GxE) (27). Here we aimed to test if GEAs in putative *cis*-regulatory contexts could predict GxE for drought better than other parts of the genome. We used published data on 176 accessions in our resequencing dataset (including only 2 African landraces) that were tested at two sites across two years in eastern Brazil, under well-watered and drought treatments (91). We used variants within ACRs, UMRs, and proximal TEs to generate growing season length genetic scores for these accessions, tested how well these scores predicted grain-yield response to drought (Pearson’s r). We then compared this observed prediction accuracy with a null distribution from matched SNPs (Fig. 5; SI Appendix, Table S5).

**Figure 5.**
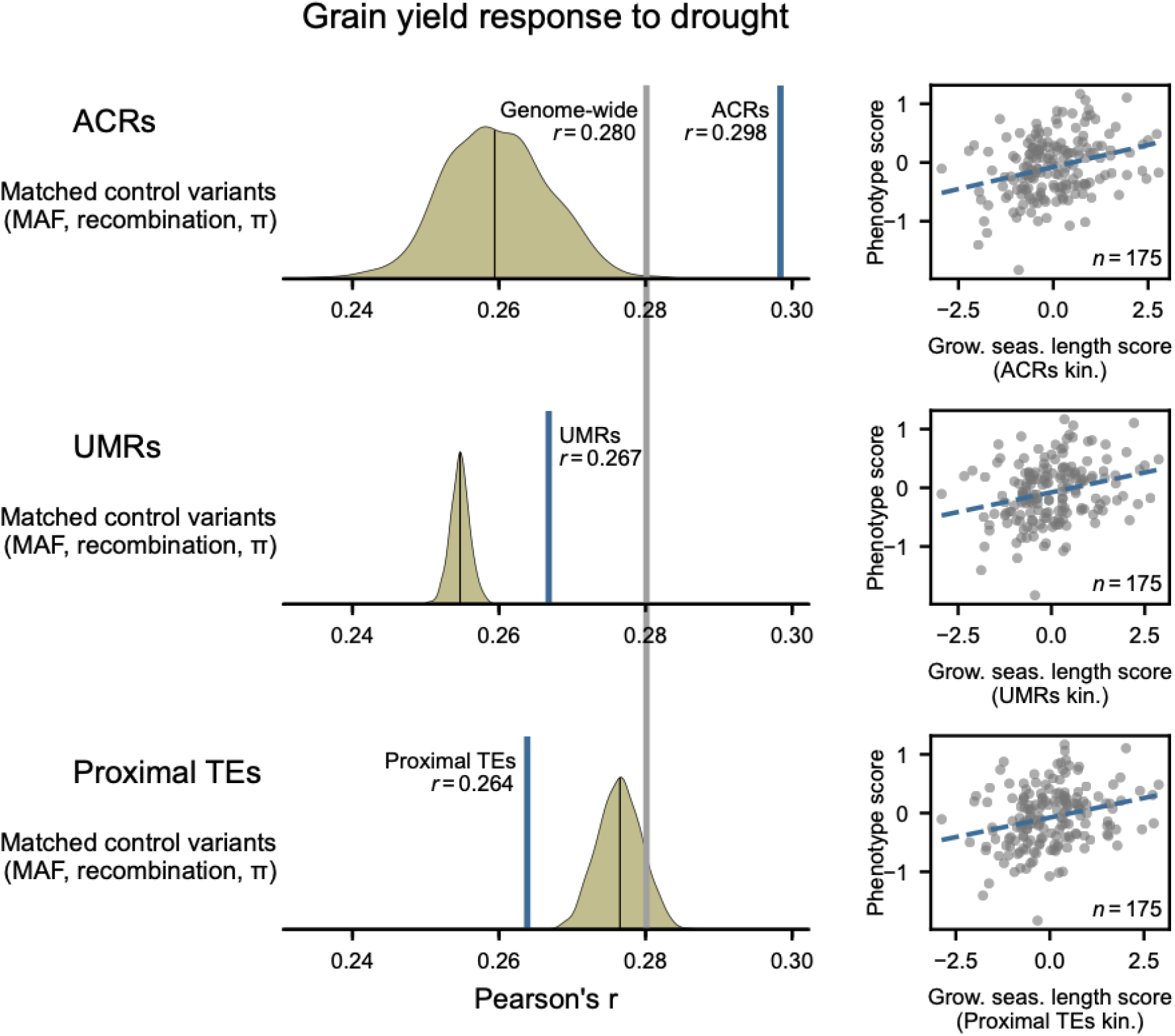
GEAs for sites in ACRs and UMRs are better predictors of genetic variation in grain yield response to drought, compared to matched control variants. Growing-season length GEAs were used to predict grain yield response to drought using genomic relationship matrices constructed from genotypes at sites within each genomic context. For each context, the left panel shows the prediction accuracy, measured as Pearson’s r, compared with matched-control site sets. Blue vertical lines indicate prediction accuracy for sites within the focal genomic context, grey vertical lines indicate prediction accuracy using genome-wide sites, and beige distributions show null expectations from 1,000 randomly sampled site sets matched to the focal context by minor allele frequency, recombination rate, and nucleotide diversity (π). Right panels show observed phenotype scores against GEA predictions from the corresponding context-specific GRM. Blue dashed lines indicate linear regression fits.

We found that growing season length-associated variants in ACRs predicted grain yield responses to drought significantly better than both matched control sites and genome-wide sites (*r*=0.292 vs. 0.271 for matched controls, *P*=0.003, *z*=3.0; genome-wide *r*=0.280), supporting the idea that drought-adaptive genetic variation is concentrated in ACRs. UMR GEAs also predicted drought response significantly better than randomly matched control sites, but less well than genome-wide sites (*r*=0.276 vs. 0.268 for matched controls, *P*≤0.001, *z*=5.9), suggesting that the frequency, diversity, and recombination rates (i.e. variables used to match) in UMRs constrain their predictive ability. In contrast, GEAs in proximal TEs predicted grain yield responses to drought less than both matched control sites and genome-wide sites (*r*=0.275 vs. 0.284 for matched controls, *P*=0.004, *z*=−2.8), despite the significant enrichment of proximal TEs in GEA outliers found above. This discrepancy may indicate that GEA enrichment in proximal TEs is driven by a modest number of strong outlier loci or partly reflects linkage disequilibrium with nearby functional sites. Collectively, these results show that signatures of local adaptation in some *cis-*regulatory loci can predict drought adaptation phenotypes.

## Discussion

Genome sequencing of genebanks yields a large number of variants that could be rapidly used in crop improvement (32, 92, 93), while traditional approaches to mining genebank diversity involve costly and difficult phenotypic screening (8, 94). Here, we used multiple lines of evidence of the genetic basis of adaptation to identify functional genetic contexts most overrepresented in loci contributing to local adaptation to climate, soil, and parasites in the globally important cereal sorghum.

Previous studies have found individual loci with diverse molecular mechanisms contributing to local adaptation including *cis*-regulatory variants (95) and amino acid substitutions in maize landraces (96), and loss of function variants in sorghum landraces and Arabidopsis (34, 62, 63). Genomic studies in diverse species have shown that *cis*-regulatory variants comprise a large fraction of QTL (97, 98), distinct ecotypes show genome-wide divergence in *cis*-regulatory variants (99–101), and GEAs are sometimes enriched in genic (58) or potential *cis*-regulatory regions (102, 103). However, there have been few attempts to study the genome-wide signal of local adaptation in crop landraces across diverse genomic contexts.

We compared a range of functional genetic assays and predictions and found the strongest and most consistent signals in likely *cis*-regulatory sequences, followed by genic sequences. In particular, sites around the transcription start site and in accessible chromatin and unmethylated regions showed the strongest and most consistent enrichment. These are likely hotspots of sequence motifs and mutations that influence transcription factor binding (18). While transcription start sites are relatively straightforward to identify from genome assemblies, more distal regulatory regions may require additional data from additional assays (104, 105). For example, assays like whole-genome bisulphite sequencing or alternatively Oxford Nanopore sequencing (106) to identify UMRs important for transcription factor binding could be useful to identify the broad set of loci potentially important in regulation and adaptation. Crops with very large genomes (e.g. *Pinus, Allium*) face substantial genotyping hurdles due to the vastness of non-coding sequence; assays capable of revealing regulatory sequences suggest a route for efficient genotyping of these sites with targeted capture or arrays. While putative *cis*-regulatory sequences were the strongest signal, we also found significant enrichment in genic regions, suggesting that changes in the amino acid sequence of proteins frequently are involved in local adaptation. Thus, our findings suggest a hierarchy of priority for experimental follow-up of potential locally adapted variants, from likely *cis-*regulatory sites to gene sequences.

A remaining goal is to discover the functional genetic basis of local adaptation in other species with different mating systems and life history, i.e. factors influencing genetic architecture. For example, local adaptation in self-pollinating crops like sorghum might have a distinct architecture compared to outcrossers like maize or pearl millet (107, 108). The architecture of local adaptation may also be influenced by gene flow between environments (14) and range expansion, which may differ substantially between regions of widespread crops (109, 110). Nevertheless, the apparent importance of *cis-*regulatory variants in sorghum is consistent with findings in outcrossing maize that QTL for putatively important phenotypes tend to be enriched in *cis-*regulatory contexts (19, 111). A priority for future work is to determine the effects of different functional types of mutations on multiple traits and fitness to determine their pleiotropy and distribution of effect sizes, which would enable synthesis of molecular, population and quantitative genetic ideas about adaptation.

### Conclusion

Beyond breeding, evolutionary biologists in other systems may consider that *cis-*regulatory variants could be among the most important mechanisms of population adaptation to environments past, present, and future. A barrier for crop researchers pursuing *cis*-regulatory variants has been there was little information available for identifying these from the many non-functional noncoding variants. The advent of molecular and computational tools like those leveraged here is rapidly unveiling these previously murky dimensions of genomes (104).

Genebanks are immense stores of natural variants, but discovering those adapted to target environments is limited by experimental logistics. Researchers working from reverse (e.g. from homology with cloned genes) or forward (e.g. from GWAS) genetics candidate genes could prioritize variants by their functional context and patterns of natural variation (9). Our findings suggest that crop geneticists and breeders should not overlook putative QTL in non-coding sequences. Instead, non-coding variants in likely *cis-*regulatory contexts should be perhaps weighted more heavily than coding sequence variants.

## Acknowledgments

Funding was provided by Gates Foundation grant INV-030574 to GPM and JRL and NIH grant R35GM138300 to JRL.

## Methods

### Data

We used the sorghum whole-genome resequencing dataset reported by Morris et al. (32), which includes 1,988 accessions sampled worldwide. In that study, reads were aligned to the *Sorghum bicolor* BTx623 v5.1 reference genome and variants were called genome-wide. We directly used the published SNP and indel calls for genotype-based analyses. GEA analyses were conducted in 443 African landraces with georeferenced collection sites. *cis*-eQTL mapping and genomic prediction used accessions from the same resequencing panel for which matched expression or phenotypic data were available (Dataset S1).

To characterize the environments experienced at landrace origins, we compiled 31 environmental variables representing climate, water availability, soil properties, radiation, topography, and biotic stress, including a modeled *Striga* HS score (SI Appendix, Table S1). Drought-related phenotypes were obtained from the Brazil field trials reported by Bernardino et al. (91), retaining 176 accessions that could be matched to the resequencing panel (Dataset S1 and Dataset S5). *cis*-eQTL analyses used the leaf RNA-seq dataset reported by Mangal et al. (112), in which 318 accessions had both genotype and expression data after matching to the resequencing panel (Dataset S1). Gene coexpression analyses used uniformly processed public sorghum RNA-seq datasets from the NCBI Sequence Read Archive (Dataset S6).

### Genomic contexts

Site- and gene-level annotations were based on the Sorghum bicolor BTx623 v5.1 reference genome. At the site level, we annotated genomic structure, epigenomic features, conservation, predicted RNA structural effects, allele frequency, nucleotide diversity, and recombination rate. Sites were classified as genic regions, proximal TEs, distal TEs, simple repeats, or other intergenic regions. Genic regions included gene bodies and 2-kb flanking sequences. Proximal and distal TEs were defined as TEs within or beyond 5 kb of gene bodies, respectively. Gene-associated variants were further assigned to upstream, UTR, coding sequence, intron, or downstream regions. Coding variants were classified as synonymous, missense, or putative loss-of-function mutations, with putative loss-of-function mutations including frameshift mutations, premature stop codons, and loss of start codons (SI Appendix, Table S2).

Epigenomic features were defined using published sorghum leaf ACRs and UMRs, which were further separated into genic and distal classes (SI Appendix, Table S2). Putative riboSNitches were annotated as variants predicted to alter local RNA secondary structure (SI Appendix, Table S2). Phylogenetic conservation was summarized using phyloP scores (113). MAF was calculated from the 443 landraces used for GEA, and nucleotide diversity and recombination rate were estimated in genomic windows and binned by genome-wide quantiles (SI Appendix, Table S2). At the gene level, we annotated expression pattern, coexpression network position, recombination environment, and gene family history. Public RNA-seq datasets were used to classify genes by expression pattern and coexpression network position. Grass orthogroups were used to classify sorghum genes as single-copy, multi-copy, or orphan genes (SI Appendix, Table S3).

### Association analyses

We used mixed linear models implemented in GEMMA v0.98.3 (114) for envGWAS and *cis*-eQTL analyses. Kinship matrices were calculated separately for each matched genotype subset and included in the models to control for population structure and relatedness among accessions. For environmental association analyses, each environmental variable was tested separately in the 443 georeferenced African landraces. We retained variants with MAF ≥ 0.05 and heterozygous genotype frequency ≤ 5%. We then aggregated site-level envGWAS results to gene-level statistics using MAGMA v1.10 (115), defining genes as gene bodies plus 2-kb flanking regions. For *cis*-eQTL mapping, we tested expressed genes in 318 accessions with matched genotype and leaf expression data. Candidate *cis* variants were defined as variants located within gene bodies or 15 kb upstream or downstream of each gene. To control for multiple testing, we used an empirical FDR framework based on the minimum *P* value within each *cis* window in observed and permuted expression datasets. SNP–gene pairs passing FDR ≤ 0.05 were called significant *cis*-eQTLs (Dataset S8).

### Enrichment analyses

To assess whether the GEA framework recovered known flowering-time biology, we first tested enrichment of GEA signals in 312 sorghum flowering-time genes mapped from FLOR-ID (40) Arabidopsis genes using orthology assignments (Dataset S4). This analysis used the same permutation-based gene-level enrichment framework described below and was performed for the four focal environmental variables. To ask whether the growing-season-length signal in flowering-time genes could reflect *cis*-regulatory variation, we further defined a conservative set of 52 canonical sorghum flowering-time genes from conserved Arabidopsis and rice flowering regulators. For genes with significant *cis*-eQTLs, the lead *cis*-eQTL was compared with the strongest top 1% growing-season-length envGWAS variant in the same *cis* interval, and genes with LD |r|>0.5 between the two variants were defined as possible *cis*-regulatory adaptation genes. Enrichment of the 52 canonical flowering-time genes among these genes was tested using a one-sided hypergeometric test, with all genes carrying significant *cis*-eQTLs as the background (Dataset S4).

For genomic-context enrichment, we ranked variants by envGWAS *P* value for each environmental variable and partitioned them by site-level context. We evaluated enrichment among the top 5% and top 1% associated sites. The top 5% threshold was used for main-text figures, and both thresholds were reported in SI Appendix, Table S2. Significance was assessed by shifting genomic context labels along the genome over 1,000 permutations, preserving relative annotation structure and local linkage disequilibrium. Observed values were standardized against empirical null distributions to obtain enrichment z scores. For gene-level enrichment, we ranked MAGMA gene-level envGWAS results by association significance and compared context-defined gene sets with the top 5% or top 1% genes. Null distributions were generated from 1,000 random gene sets of equal size (SI Appendix, Table S3).

### Variance partitioning

For each focal genomic context, we estimated the proportion of environmental variation explained by genotypes at sites in that context. Analyses focused on ACRs, UMRs, and proximal TEs. For each context, we compared the observed variance explained with two references: a genome-wide estimate based on all eligible sites, and an empirical null distribution generated from random site sets matched for site number, MAF, recombination rate, and nucleotide diversity. Variance components were estimated by REML in LDAK v5.2 (116) (SI Appendix, Table S4). Significance was assessed with 1,000 matched-site permutations. Contexts exceeding the matched random expectation were interpreted as explaining more environmental variation than expected for comparable sites.

### Genomic prediction of drought response

Growing season length scores were generated with a best linear unbiased prediction (BLUP) framework implemented in LDAK v5.2 (116) (SI Appendix, Table S5). For ACRs, UMRs, proximal TEs, and genome-wide sites, we fit growing-season-length models in the georeferenced landrace panel, estimated BLUP effects, and calculated predicted growing season length scores for accessions from the Brazil drought trials. Prediction accuracy was evaluated as the Pearson correlation between predicted growing season length scores and grain-yield response to drought, measured as grain-yield plasticity. Significance for each context was assessed with empirical null distributions generated from 1,000 random site sets matched for site number, minor allele frequency, recombination rate, and nucleotide diversity.

