## Supplemental appendix for "Functional genomic map of local adaptation in sorghum to guide allele mining"

### Supporting Information Text

#### Supplementary Note 1. Genotype data and accession subsets

We used the sorghum whole-genome resequencing dataset reported by Morris et al. (1) as the genotype basis for all analyses. This dataset includes 1,988 accessions sampled worldwide. In the original study, reads were aligned to the *Sorghum bicolor* BTx623 v5.1 reference genome, and genome-wide single-nucleotide polymorphism (SNP) and insertion-deletion (InDels) variants were called. We used the published SNP and InDels calls directly, without repeating read alignment or variant calling. All coordinate-based annotations and genomic-context assignments were performed in the BTx623 v5.1 reference coordinate system (1).

*Genotype–environment association.*

Genotype–environment association (GEA) analyses were conducted in 443 African georeferenced landraces (Dataset S1). These accessions were used for environmental variable extraction, redundancy analysis, environmental GWAS, site- and gene-level enrichment analyses based on environmental association signals, and variance partitioning of environmental variables by genomic context, as described below.

*cis-eQTL analyses.*

*cis*-eQTL analyses used accessions that overlapped between the Morris et al. (1) resequencing panel and the sorghum leaf RNA-seq dataset reported by Mangal et al. (2). After matching accession identifiers between the two datasets, 318 accessions had both genotype and leaf expression data and were retained for *cis*-eQTL mapping (Dataset S1).

*Genomic prediction of drought response.*

Genomic prediction analyses used accessions that overlapped between the resequencing panel and the Brazil drought field trials reported by Bernardino et al. (3). A total of 176 accessions with matched genotype and drought-related phenotype data were retained for prediction analyses (Dataset S1).

#### Supplementary Note 2. Environmental variables and selection of focal environmental gradients

To characterize the environments experienced at landrace origins, we compiled 31 environmental variables (Table S1) following the general framework used in earlier sorghum local adaptation studies (4). These variables represent major ecological axes, including climate, water availability, soil properties, radiation, topography, and biotic stress. All variables were extracted from gridded environmental layers at accession collection coordinates using global- or Africa-scale raster datasets.

*Climate and water availability.*

Climate was characterized primarily with bioclimatic variables from CHELSA v2.1 (5), which summarize long-term temperature and precipitation regimes, including means, seasonality, and extremes. We included multiple temperature and precipitation variables, together with the aridity index, to capture water balance and drought gradients (6). We also included growing season length, which reflects climatic effects on the duration of the potential crop growing period (5).

*Radiation and topography.*

To capture seasonal radiation environments, we calculated mean quarterly photosynthetically active radiation from the NASA Surface Radiation Budget dataset (https://asdc.larc.nasa.gov/project/SRB). Topography was represented by altitude and topographic wetness index. Altitude captures broad elevational gradients associated with temperature and precipitation, whereas TWI reflects topographic effects on surface water accumulation and soil moisture distribution (7).

*Soil and hydrological variables.*

To represent soil chemistry, nutrient status, and water-holding capacity, we included topsoil pH, extractable aluminum, and total nitrogen and phosphorus in the 0–30 cm soil layer (8). To capture soil physical properties related to water retention, we also included clay content and extractable soil moisture capacity at 5 cm depth (9). In addition, depth to the water table was included as an index of potential groundwater supply (10).

*Striga prevalence.*

Because *Striga hermonthica* is a major biotic constraint on sorghum production in Africa, we developed a *Striga* habitat suitability (HS) score to estimate potential parasite pressure at each collection site. Species distribution modeling generally followed Bellis et al. (11), but with an expanded occurrence dataset that included 1,369 records compiled by Bellis et al., 384 additional field observations from this study, and 1,762 records downloaded from GBIF. To reduce spatial sampling bias, occurrence records were spatially filtered to remove duplicates within 0.01° (~1 km), leaving 1,334 occurrence points for model training (Dataset S2). We used MaxEnt v3.4.4 (12) to predict potential suitability from relationships between *Striga* occurrence and environmental variables. Model tuning and evaluation were performed with ENMeval v2.0.0 (13) using the "checkerboard2" partitioning scheme to reduce the effect of spatial autocorrelation. Among candidate models, we selected the model with the lowest corrected Akaike information criterion (AICc) and extracted predicted HS values at sorghum collection coordinates. To avoid overestimating parasite pressure in areas lacking parasite records, accessions located >200 km from any known *S. hermonthica* record were assigned HS = 0. We generated two versions of this score. The first used the same environmental predictor classes as Bellis et al. (11), including climate, topography, and soil variables (Dataset S3), and is referred to as the *Striga* HS score. The second additionally incorporates crop distribution data and is referred to as the *Striga* HS score (Crop).

*Redundancy Analysis.*

To assess whether the 443 African sorghum landraces spanned broad variation in the 31 environmental variables and whether these variables explained genome-wide genotypic variation, we used redundancy analysis (RDA) to quantify the fraction of genome-wide genotypic variation explained by linear combinations of the environmental variables (14). RDA was performed using environmental variables as predictors and genome-wide SNP genotypes as multivariate responses. Environmental variables were standardized, landraces with missing environmental values were excluded, and the genotype matrix was centered before analysis. The centered genotype matrix was then regressed on the standardized environmental matrix, and principal component decomposition was applied to the fitted genotype values to obtain constrained axes and their variance contributions. Overall explained variance was calculated by comparing fitted genotypic variation with total centered genotypic variation, with adjusted $R^2$ used to account for the number of predictors and samples. For the RDA biplot (Fig. 1C), landraces were plotted using their scores on the first two constrained axes, and environmental-variable vectors were drawn from the correlations between standardized environmental variables and RDA sample scores. Vector direction indicates the environmental variables associated with each axis, and vector length reflects the strength of the association. Analyses were implemented in Python v3.9.19 using scikit-learn v1.5.2 (15).

Across 987,993 unlinked SNPs ($r^2 < 0.5$), the 31 environmental variables explained 18.7% of genome-wide genotypic variation. The first constrained axis explained 4.5% of genome-wide genotypic variation and was associated primarily with temperature and elevation variables, together with some correlated soil variables. The second constrained axis explained 3.6% of genome-wide genotypic variation and was associated mainly with precipitation and radiation variables (Fig. 1C). These results indicate that environmental variables captured substantial structure in genome-wide genotypic variation among African sorghum landraces.

*Environmental correlation structure and focal-variable selection.*

The RDA biplot showed that many environmental variables were oriented in similar directions, suggesting that some predictors captured overlapping environmental gradients. We therefore directly examined correlation structure among the 31 environmental variables across the 443 African sorghum landraces. Pairwise Pearson correlations were calculated among all environmental variables and visualized as an environmental correlation network (Fig. 1D). In this network, nodes represent environmental variables, node size reflects the number of strong correlations for each variable, and edges indicate correlations with $|r| > 0.7$, with edge color showing the direction and magnitude of the correlation. This analysis confirmed substantial collinearity among several environmental variables. Temperature-related, precipitation-related, radiation-related, and soil/hydrology variables each formed correlated clusters, indicating that many variables represented overlapping aspects of the same broad environmental gradients. We used this correlation structure, together with prior evidence for sorghum adaptation to drought (4), aluminum toxicity (4), and *Striga hermonthica* pressure (11), to select four focal environmental variables for detailed downstream analyses.

Drought-related variables formed two major clusters, reflecting complementary aspects of water limitation across African environments (Fig. 1D). We selected growing season length (GSL) and precipitation of the warmest quarter (PHQ) as two drought-related gradients. GSL reflects the duration of the potential growing period and is closely related to terminal drought, whereas PHQ captures water availability during hot growing-season conditions. We further selected soil extractable aluminum as a proxy for aluminum toxicity and the *Striga* HS score as a proxy for parasite pressure. All 31 environmental variables were retained for RDA and genome-wide environmental association analyses, whereas these four focal variables were used for detailed enrichment analyses, variance partitioning, prediction analyses, and biological interpretation of genomic-context-specific signals (see below).

**Supplementary Note 3. *cis*-eQTL mapping**

*cis*-eQTL analyses used the sorghum leaf RNA-seq dataset reported by Mangal et al. (2). In that study, 738 accessions were grown in a single field experiment at the University of Nebraska–Lincoln's Havelock Research Farm in Lincoln, Nebraska. Mature leaf tissue was collected within a single day and over a sampling window of less than two hours using a standardized protocol, minimizing variation due to environment, tissue type, developmental stage, and sampling time. After matching accessions to the Morris et al. (1) resequencing panel, 318 accessions had both genotype and leaf expression data and were retained for *cis*-eQTL mapping (Dataset S1).

We obtained the Transcripts Per Million (TPM) expression matrix directly from Mangal et al. (12). Genes were retained for *cis*-eQTL mapping if they had TPM > 1 in at least 20% of the 318 accessions. Expression values for retained genes were transformed as $\log_2(\text{TPM} + 1)$ and then inverse-normal transformed across accessions. The transformed expression matrix was used as the phenotype matrix for *cis*-eQTL mapping.

Candidate *cis* variants were defined as variants located within gene bodies or within 15 kb upstream or downstream of each gene. Each variant–gene pair within this *cis* window was tested using the mixed linear model implemented in GEMMA v0.98.3 (16) with kinship control. For the 318 accessions used in *cis*-eQTL mapping, the kinship matrix was calculated in GEMMA using LD-pruned SNPs filtered for minor allele frequency (MAF) ≥ 0.01, heterozygous genotype frequency ≤ 0.01, and pairwise $r^2 < 0.5$ within 15-kb sliding windows. The resulting kinship matrix was included as the random-effect covariance matrix in the mixed model.

Multiple testing was controlled using the empirical FDR framework of Westra et al. (17). Briefly, for each gene, we recorded the minimum *P* value across all variants in its *cis* window. We then permuted expression phenotypes 100 times, repeated the *cis*-eQTL scan for each permutation, and recorded the minimum *cis*-window *P* value for each gene in each permuted dataset. Observed and permuted minimum *P* values were used to estimate empirical FDR across significance thresholds. Variant–gene pairs passing the FDR = 0.05 threshold were called significant *cis*-eQTLs. Significant *cis*-eQTLs were detected for 12,454 genes and are reported in Dataset S8.

**Supplementary Note 4. Public RNA-seq processing, expression classes, and coexpression networks**

To characterize gene expression patterns across diverse tissues, developmental stages, treatments, and stress conditions, we collected public Sorghum RNA-seq datasets from the NCBI Sequence Read Archive (SRA). We searched SRA using the keywords "*Sorghum bicolor*" and "*Sorghum bicolor subsp.*" on September 21, 2024. We restricted results to assay type RNA-seq and retained datasets with sequencing depth <20 Gb. This search yielded 7,137 RNA-seq datasets for uniform processing (Dataset S6).

*Data Preprocessing and Alignment.*

Raw reads were quality filtered with fastp v0.23.4 (18) and aligned to the *Sorghum bicolor* BTx623 v5.1 reference genome using HISAT2 v2.2.1 (19) in spliced-alignment mode. Gene-level read counts were quantified with featureCounts v1.6.2 (20). Samples were retained if they had ≥5 million clean reads, ≥80% clean read proportion, ≥70% genome alignment rate, and ≥50% of reads assigned to protein-coding genes. After filtering, 4,591 high-quality RNA-seq datasets remained. To reduce redundancy from biological replicates or highly similar samples, we performed pairwise differential expression analyses using edgeR v4.0.16 (21). Samples with fewer than 50 differentially expressed genes relative to another sample were considered redundant and removed. This procedure resulted in a final set of 1,923 RNA-seq datasets for downstream expression classification and coexpression network construction.

*Gene expression classes.*

Gene-level read counts were converted to TPM. Among 31,819 nuclear genes, genes with TPM ≤ 2 across all 1,923 samples were classified as genes with no detectable expression. Because expression variance scales with mean expression, we modeled the relationship between mean log2(TPM+1) and expression variance using locally estimated scatterplot smoothing (LOESS) implemented in the Python package statsmodels v0.14.4 (22). Genes were classified based on standardized residuals from this model. Genes with residuals > 1 were defined as high-variance genes, whereas genes with residuals < 1 and TPM > 2 in all samples were classified as stably expressed genes.

*Gene coexpression networks.*

We constructed a global gene coexpression network following Wisecaver et al. (23). Using the 1,923 RNA-seq datasets, we calculated Pearson correlation coefficients (PCCs) between all gene pairs based on TPM values and ranked coexpressed genes for each gene. For each gene pair, mutual rank (MR) was calculated as the geometric mean of their reciprocal PCC ranks. MR values were converted to edge weights using an exponential decay function, and overlapping coexpression modules were identified with ClusterONE v1.0 (24) from the resulting weighted network using parameters n = 3 and minimum edge weight = 0.1.

In total, 5,095 coexpression modules were detected (Dataset S7), covering 27,552 genes. The remaining 4,267 genes were classified as non-module genes. For modules containing at least five genes, we calculated four centrality metrics for each gene: degree, closeness, betweenness, and eigenvector centrality with the Python package NetworkX v3.0 (25). Each metric was standardized as a $z$ score within the network. Genes with $z$ scores > 1 for at least two metrics were classified as module hubs, resulting in 7,135 hub genes. The remaining 20,417 genes in modules were classified as non-hub module genes (Table S3).

**Supplementary Note 5. Site-level genomic context annotation**

We annotated sites according to genomic structure, local gene features, coding effects, epigenomic features, predicted RNA structural effects, phylogenetic conservation, allele frequency, nucleotide diversity, and recombination rate.

*Genomic structure and local gene features.*

Genomic structure was annotated from the BTx623 v5.1 gene and repeat annotations (1). Sites were classified into five broad categories according to their position relative to genes and repetitive elements: genic regions, proximal transposable elements (TEs), distal TEs, simple repeats, and other intergenic regions. Genic regions were defined as gene bodies plus 2-kb upstream and downstream flanking sequences. Proximal TEs were defined as TEs located within 5 kb of gene bodies, whereas distal TEs were defined as TEs more than 5 kb from gene bodies (Fig. 2A and Table S2). Sites overlapping multiple categories were assigned according to a predefined priority order. Within gene-associated regions, variants were further classified as upstream 2 kb, 5′ UTR, coding sequence, intron, 3′ UTR, or downstream 2 kb (Table S2).

*Coding variant effects.*

Coding variant annotations were obtained from the SnpEff-based variant effect predictions reported by Morris et al. (1). We used these annotations to classify variants in coding sequences according to their predicted effects on protein sequences. Coding variants were grouped into synonymous, missense, and putative loss-of-function mutations. Putative loss-of-function mutations included frameshift mutations, premature stop codons, and loss of start codons (Table S2).

*Epigenomic features.*

Accessible chromatin regions (ACRs) were obtained from sorghum leaf ATAC-seq data reported by Zhou et al. (26). Unmethylated regions (UMRs) were obtained from whole-genome bisulfite sequencing data reported by Crisp et al. (27). ACRs and UMRs were intersected with variant positions to define epigenomic site-level contexts. Regions located within 2 kb of gene bodies were classified as genic ACRs or genic UMRs, whereas regions farther than 2 kb from gene bodies were classified as distal ACRs or distal UMRs (Fig. 2B and Table S2).

*RiboSnitches.*

We defined putative riboSnitches as SNPs predicted to alter local RNA secondary structure. RiboSnitches were predicted with SNPfold v1.01 (28), following the genome-wide approach and correlation threshold used by Ferrero-Serrano et al. (29). SNPfold compares two equal-length allele-specific sequences that differ at a focal variant by evaluating the ensemble of possible RNA secondary structures for each sequence. It then compares their base-pairing probability matrices and returns a correlation coefficient that reflects the similarity of the predicted structural ensembles. Lower correlation coefficients therefore indicate stronger predicted allele-dependent changes in RNA secondary structure.

We restricted this analysis to SNPs within annotated gene bodies, including exons and introns. For each candidate SNP, we extracted the focal site and 40 nucleotides on each side from the BTx623 v5.1 reference genome, producing an 81-nt window centered on the SNP. The reference allele sequence was taken directly from the reference genome, and the alternate allele sequence was generated by substituting the alternate allele at the focal site. We then ran SNPfold with default parameters to compare the reference and alternate allele sequences. SNPs with a SNPfold correlation coefficient < 0.8 were classified as putative riboSnitches. SNPs were excluded if a complete 40-nt flanking sequence could not be obtained on both sides (Fig. 2E and Table S2).

*Phylogenetic conservation.*

Phylogenetic conservation scores were obtained from PlantRegMap (30), which generated genome-wide conservation landscapes from multispecies plant genome alignments and computed base-wise conservation scores with PHAST. We used phyloP scores, which measure conservation or acceleration relative to a neutral model. For sorghum, scores were calculated within the PACMAD clade of Poales, a major clade of grasses (Poaceae), spanning ~33 million years of evolution and including tropical cereals like sorghum, maize, pearl millet, and teff (31). Downstream analyses were restricted to sites within genes and 15-kb upstream and downstream flanking regions. Regions lacking sufficient phylogenetic information for conservation inference, because of limited alignment coverage, repeat-associated masking, or weak phylogenetic signal, were classified as non-evaluable. Among evaluable sites, phyloP > 1 was classified as conserved, phyloP < −1 was classified as accelerated, and the remaining sites were classified as neutral (Fig. S7 and Table S2).

*Population genetic context.*

Population genetic analyses were restricted to sites within genes and 15-kb upstream and downstream flanking regions. MAF was calculated from genotype data for the 443 African landraces used for GEA. Variants were grouped into five MAF bins: 0.05–0.10, 0.10–0.20, 0.20–0.30, 0.30–0.40, and 0.40–0.50. Nucleotide diversity ($\pi$) was calculated with scikit-allel v1.3.8 (32) in 500-bp windows and binned by genome-wide quantiles into high-diversity regions, intermediate-diversity regions, and low-diversity regions. High-diversity regions were defined as the top 25% of windows, and low-diversity regions were defined as the bottom 25% (Fig. S7 and Table S2).

**Supplementary Note 6. Gene-level genomic context annotation**

Gene-level genomic contexts were assigned from expression pattern, coexpression network position, recombination environment, and gene family history. All gene-level annotations were based on BTx623 v5.1 gene models. Expression classes and Coexpression network position see Supplementary Note 4.

*Recombination context.*

Genome-wide recombination rates were estimated from family-level genetic maps inferred from the sorghum NAM population (1). Briefly, genetic maps were constructed on the *Sorghum bicolor* v5.1 reference genome using custom scripts, and recombination rates were calculated in sliding windows and averaged across families to generate a genome-wide recombination landscape. We summarized recombination rates in 1-kb windows and binned windows by genome-wide quantiles into high-, intermediate-, and low-recombination regions.

High-recombination regions were defined as the top 25% of windows, and low-recombination regions were defined as the bottom 25%. Genes were assigned to recombination categories according to the windows in which they were located (Fig. S7 and Table S3).

*Gene family history.*

Gene family history was inferred from the protein sequences of annotated gene models from six grasses (*Sorghum bicolor*, *Zea mays*, *Oropetium thomaeum*, *Brachypodium distachyon*, *Oryza sativa*, and *Pharus latifolius*) together with several outgroup species. Orthogroups were inferred with OrthoFinder v2.5.5 (33) using default parameters. Protein sequences within each orthogroup were aligned with MAFFT v7.526 (34), and gene trees were reconstructed with FastTree v2.1.11 (35). From these results, we defined grass-level hierarchical orthogroups at the most recent common ancestor of grasses, corresponding approximately to the Poaceae common ancestor (~69 MYA) (31). Sorghum genes assigned to these grass-level orthogroups were classified as single-copy or multi-copy according to their copy number within each orthogroup. Genes not assigned to any grass-level orthogroup were classified as orphan genes, which may include potential de novo genes, rapidly diverged sequences, or horizontally transferred genes (Table S3).

**Supplementary Note 7. Environmental GWAS and gene-level aggregation**

Environmental association analyses were conducted using mixed linear models implemented in GEMMA v0.98.3 (16). Each environmental variable was tested separately in the 443 georeferenced African landraces. To ensure stable inference, we retained variants with MAF ≥ 0.05 and heterozygous genotype frequency ≤ 5%.

Kinship matrices were calculated separately for each analysis using the corresponding genotype subset. SNPs used for kinship estimation were first pruned for linkage disequilibrium in 15-kb sliding windows using Pearson's $r^2 < 0.5$. Additional filters retained variants with MAF ≥ 0.01 and heterozygous genotype frequency ≤ 0.01. Identity-by-state distances were then calculated from the filtered SNPs and used to construct kinship matrices. These kinship matrices were included in the GEMMA mixed linear models to control for population structure and relatedness among accessions.

Site-level envGWAS results were aggregated to gene-level statistics using MAGMA v1.10 (36). Gene regions were defined as gene bodies plus 2-kb upstream and downstream flanking sequences. For each environmental variable, variant-level association statistics were mapped to genes and combined using MAGMA to obtain gene-level *P* values. These gene-level envGWAS results were used for gene ranking, gene-level enrichment analyses, flowering-time gene enrichment tests, and sliding-window analyses around candidate genes.

**Supplementary Note 8. Flowering-time gene sets and *cis*-regulatory adaptation genes**

*Flowering-time gene enrichment in GEA signals*

We first asked whether gene-level GEA signals were enriched for known flowering-time genes. To define a broad sorghum flowering-time gene set, we mapped Arabidopsis flowering-time genes curated in FLOR-ID (37) to their sorghum orthologs, identifying 312 genes (Dataset S4). We tested enrichment of these genes among GEA-associated genes using the gene-level enrichment framework described in Supplementary Note 9. GEA-associated genes were defined at two rank-based thresholds, corresponding to the top 5% and top 1% most strongly associated genes (Table S3). Flowering-time genes were enriched among genes most strongly associated with growing season length (see Main Text).

*Flowering-time gene enrichment among cis-regulatory adaptation genes*

We next asked whether the growing-season-length GEA signal in flowering-time genes might be explained, at least in part, by *cis*-regulatory variation. For this analysis, we used a more conservative set of canonical flowering-time genes. We curated conserved flowering-time regulators from Arabidopsis and rice, then identified their sorghum homologs using Phytozome orthology assignments. This yielded 52 canonical sorghum flowering-time genes (Dataset S4).

We defined possible *cis*-regulatory adaptation genes by integrating *cis*-eQTL results (see Supplementary Note 3) with growing-season-length envGWAS signals. For each gene with significant *cis*-eQTLs, we first identified the lead *cis*-eQTL, defined as the *cis* variant most strongly associated with expression. We then searched the same *cis* interval for the strongest growing-season-length envGWAS variant, restricting this search to variants in the top 1% genome-wide for growing-season-length association. Finally, we calculated linkage disequilibrium between the lead *cis*-eQTL and this envGWAS variant. Genes were classified as possible *cis*-regulatory adaptation genes when the LD correlation satisfied $|r| > 0.5$. This criterion identified 968 genes (Dataset S4).

Among the 52 canonical flowering-time genes, 25 had significant *cis*-eQTLs. Five of these were classified as possible *cis*-regulatory adaptation genes (Dataset S4). We tested this overlap with a one-sided hypergeometric test, using all genes with significant *cis*-eQTLs as the background (N=12,454, Dataset S8). Canonical flowering-time genes were significantly enriched among possible *cis*-regulatory adaptation genes ($P = 0.0405$).

**Supplementary Note 9. Enrichment analyses**

We tested whether environmental association signals were enriched or depleted across the site- and gene-level genomic contexts defined above. Analyses were performed separately for each environmental variable. Significance was assessed using permutation-based empirical null distributions.

*Site-level enrichment.*

Site-level enrichment was based on variant-level envGWAS results (see Supplementary Note 7). For each environmental variable, variants were ranked by association *P* value and, for each site-level genomic context (see Supplementary Note 5), partitioned into those inside or outside the focal context. Enrichment was quantified as the ratio of the $-\log10(P)$ value at the 5% threshold inside versus outside the context, and the same analysis was repeated at the 1% threshold (Table S2). Statistical significance was assessed using a permutation-based empirical null distribution generated by shifting genomic context labels by a random distance along the genome and recalculating the same statistic (1,000 permutations). Because this procedure preserves the relative genomic ordering of contexts, local linkage disequilibrium structure was retained. Observed values were standardized relative to the null distribution to obtain enrichment *z* scores, and two-sided empirical *P* values were calculated accordingly (Table S2).

Site-level enrichment results for the major genomic and functional contexts, including genic regions, transposable-element contexts, coding effects, ACRs, UMRs, and putative riboSnitches, are described in the Main Text and supporting figures. We also evaluated population-genetic and evolutionary contexts, including MAF, nucleotide diversity, and phylogenetic conservation, as additional site-level contexts (Table S2).

GEA enrichment across MAF and nucleotide-diversity classes differed among environmental gradients. On one hand were the drought variables, where GEAs were enriched in uncommon (e.g. 0.05 < MAF < 0.1) variants and depleted in common (e.g. MAF > 0.3) variants (uncommon variants: $z$ = 1.6 – 6.2, $P$ = 0.001 – 0.061; common variants: $z$ = −5.8 to −0.5, $P$ = 0.001 – 0.305). Additionally, aluminum-associated variants were enriched in uncommon variants (0.05 < MAF < 0.1; $z$ = 4.5, $P$ < 0.001). On the other hand were *Striga*-associated variants, which were enriched in high frequency (MAF > 0.4; $z$ = 3.8, $P$ < 0.001), high π ($z$ = 3.4, $P$ = 0.001), and depleted in low frequency (0.05 < MAF < 0.1; $z$ = -6.1, $P$ < 0.001) and low to medium π variants ($z$ = -3.4 to −3.0, $P$ ≤ 0.002; Table S2).

Phylogenetic conservation was evaluated for variants within genes and their flanking 15-kb regions. Regions lacking sufficient phylogenetic information for conservation inference (non-evaluable in Fig. S8) showed an overall depletion of GEAs across environmental variables (significantly for extractable aluminum, $z$ = −1.9, $P$ = 0.025), suggesting that cross-species alignable, non-repetitive regions near genes are more likely to contribute to local adaptation. Within these evaluable sites, we further classified variants into accelerated, neutral, and conserved categories based on phyloP scores. Aside from a slight enrichment at neutral sites (significantly for growing season length, $z$ = 2.0, $P$ = 0.019), variants with accelerated or conserved patterns of evolution were not significantly enriched for GEAs (Fig. 3F and Table S2). These results suggest that relationships between phylogenetic conservation and local adaptation signals may be weak, context-dependent, or partly obscured by opposing evolutionary processes.

*Gene-level enrichment.*

Gene-level enrichment was based on MAGMA gene-level envGWAS results (see Supplementary Note 7). For each environmental variable, genes were ranked by gene-level association significance. For each gene-level genomic context, enrichment was evaluated as the overlap between the context-defined gene set and the top 5% or top 1% most strongly associated genes. Enrichment was quantified using the Jaccard index between the context-defined gene set and the top-ranked gene set. Statistical significance was assessed using an empirical null distribution generated by randomly sampling gene sets of equal size and recalculating the same statistic over 1,000 permutations. Observed values were standardized relative to the null distribution to obtain enrichment z scores, and two-sided empirical *P* values were calculated accordingly.

Gene-level enrichment results for the major biological and functional gene categories, including flowering-time gene sets, expression classes, coexpression-network categories, and gene-family-history classes, are described in the Main Text (Fig. 3). We also evaluated the recombination environment as an additional gene-level context (Fig. S7). Genes in low-recombination regions showed a tendency toward depletion for GEA signals, although this pattern was not statistically significant across the focal environmental variables (Fig. S8 and Table S3).

*Relative enrichment around genes*:

To characterize the spatial distribution of local adaptation signals around genes, we performed sliding-window analyses around the top 5% most significant genes from the gene-level analysis. For each environmental variable, regions extending 10 kb upstream and downstream of both the transcription start site (TSS) and transcription termination site (TTS) were extracted and analyzed using 500-bp windows with a 10-bp step size. Within each window, local association strength was summarized as the first quartile (Q1) of variant $-\log10(P)$ values, and local relative signal intensity was calculated by comparing the window-specific Q1 with the overall Q1 across the corresponding ±10 kb region. Statistical significance was assessed using a permutation-based empirical null distribution generated by randomly sampling gene sets of equal size and repeating the same sliding-window analysis (1,000 permutations). The resulting permutation distributions at each window were used to derive empirical 95% intervals for local relative signal intensity, and observed values above these intervals were interpreted as enrichment, whereas values below them were interpreted as depletion.

To compare local adaptation signals with other genomic properties around genes, we calculated additional tracks in the same 500-bp windows. These included nucleotide diversity ($\pi$), ACR coverage, and UMR coverage (see Supplementary Note 5). We also calculated a high-effect index from promoter mutagenesis data reported by Groover et al. (38). That study used massively parallel reporter assays to measure expression effects of promoter and 5′ UTR mutations in three sorghum genes, PsbS, Raf1, and SBPase. We inferred single-mutation effects from barcode-level data using an additive linear model, because many constructs carried multiple mutations. Within each gene, mutations in the top 5% of absolute inferred effect size were classified as high-effect mutations. For each TSS-centered window, we then calculated the probability that mutations in that window were high-effect mutations and divided this value by the gene-weighted background probability. The resulting high-effect index measures local enrichment of experimentally observed high-effect promoter mutations, with values above 1 indicating enrichment and values below 1 indicating depletion.

**Supplementary Note 10. Variance partitioning by genomic context**

We used variance partitioning to estimate how much variation in environmental variables could be explained by genomic relationship matrices (GRMs) built from variants in specific genomic contexts. Analyses focused on ACRs, UMRs, and proximal TEs. For each context, all eligible variants in that context were used to construct a context-specific GRM with LDAK v5.2 (39). We also constructed a genome-wide GRM from all eligible variants as a common reference.

To generate a matched null expectation, we sampled 1,000 random variant sets for each context. Each random set contained the same number of variants as the focal context and was matched to the observed variants by MAF, recombination rate, and nucleotide diversity. A GRM was constructed from each matched set using the same LDAK procedure.

Variance components were estimated by restricted maximum likelihood (REML) in LDAK v5.2 (39) for the four focal environmental variables: growing season length, precipitation of the warmest quarter, extractable aluminum, and *Striga* HS score. For each variable, the context-specific, matched random, and genome-wide GRMs were fitted separately to estimate the proportion of environmental variation explained. Statistical significance was assessed by comparing the observed variance explained by each context-specific GRM with the empirical null distribution from the matched random GRMs. The genome-wide GRM was used as an additional reference, but not as the permutation-based null (Fig. 4B and Table S4).

**Supplementary Note 11. Genomic prediction of drought response**

Drought-related phenotypes were obtained from the multi-environment field trials reported by Bernardino et al. (3). That study evaluated drought response in a diverse set of sorghum accessions from the Sorghum Association Panel (SAP) in two tropical agricultural regions in Brazil, Janaúba, Minas Gerais, and Teresina, Piauí. Trials were conducted in 2010 and 2012, with Janaúba evaluated in both years and Teresina evaluated in 2010. Each site-year environment included well-watered (WW) and water-stressed (WS) treatments, allowing genotype performance to be compared under control and drought conditions.

We retained accessions that could be matched to the Morris et al. (1) resequencing panel and had complete measurements across the selected site-year environments. A total of 176 accessions had matched genotype and drought-related phenotype data and were used in prediction analyses (Dataset S5). The measured traits included grain yield, flowering time, and plant height. For each trait and site-year environment, we calculated two derived phenotypes: the average trait value across treatments, defined as (WW + WS) / 2, and drought-response plasticity, defined as log2(WW / WS). Zero values were treated as missing before ratio calculation. Derived phenotypes were standardized across accessions using z scores. Values with absolute z scores greater than 3 were treated as missing to reduce the influence of extreme observations. For each trait, standardized values were then averaged across the available site-year environments to obtain overall trait means and plasticity measures. Genomic prediction analyses focused on grain-yield plasticity, calculated as the mean standardized log2(WW / WS) grain-yield response across the Brazil field trials.

We used a best linear unbiased prediction (BLUP) framework implemented in LDAK v5.2 (39) to test whether genomic variation associated with growing season length predicted grain-yield response to drought in the Brazil trials. Prediction analyses used the same context-specific, matched-random, and genome-wide variant-set framework described in Supplementary Note 10. For each variant set, growing season length was modeled in the georeferenced landrace panel using REML, and BLUP effects from this model were used to calculate predicted growing season length scores for the Brazil trial accessions. Prediction accuracy was evaluated as the Pearson correlation between predicted growing season length scores and grain-yield plasticity. For each focal context, significance was assessed against empirical null distributions from 1,000 matched random variant sets. The genome-wide prediction score was used as an additional reference (Table S5).

### Figures

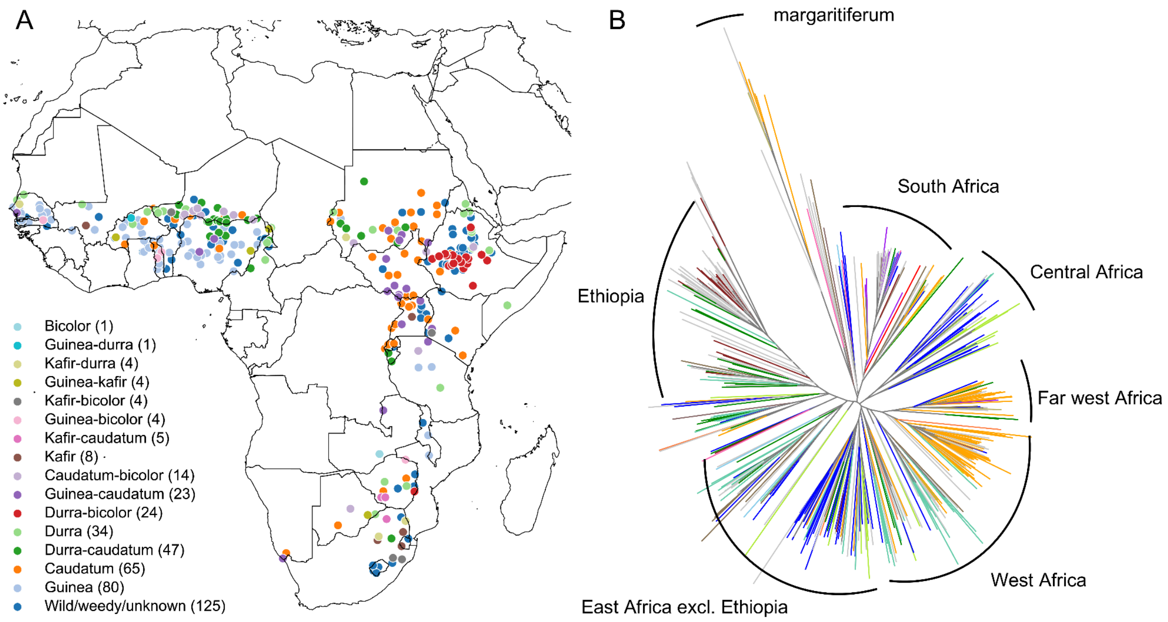

**Fig. S1. African sorghum landraces span a wide range of environments.** (A) Map showing the geographic origins of sorghum landraces analyzed in this study, with colors indicating botanical types. (B) Neighbor-joining tree constructed from LD-pruned SNPs, with major clades annotated by their predominant geographic regions.

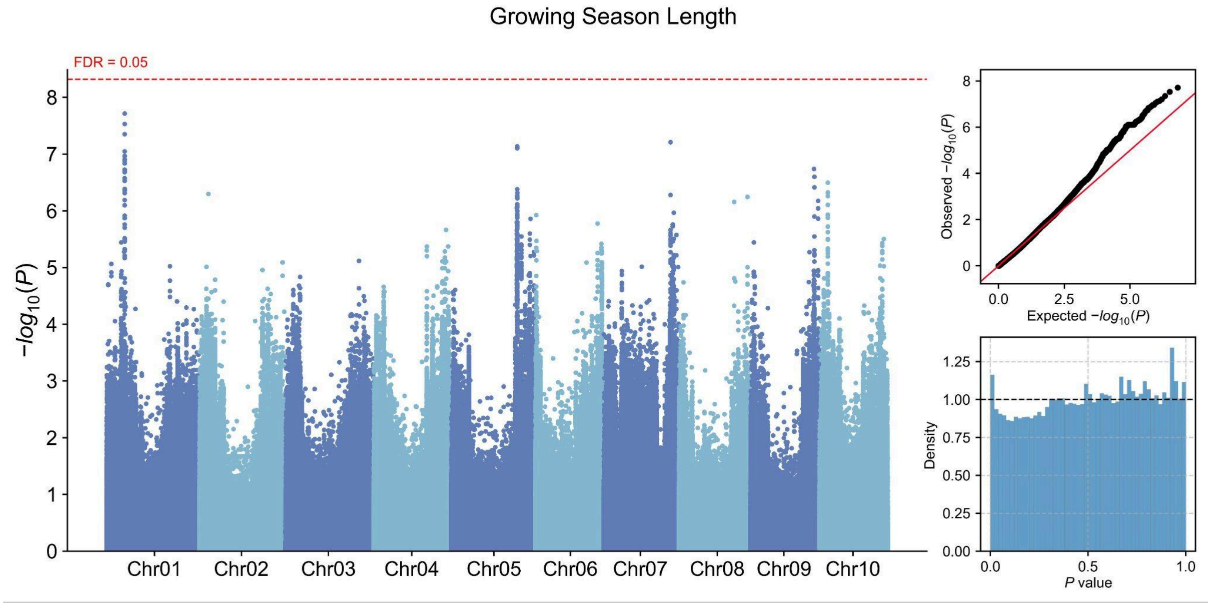

**Fig. S2. Genome-wide association results for growing season length.** The Manhattan plot shows variant associations across the genome, with each point representing a tested variant plotted by genomic position and −log10(*P*). Chromosomes are displayed in alternating colors, and the dashed red line denotes the false discovery rate (FDR) threshold of 0.05. The quantile–quantile plot compares observed and expected −log10(*P*) values under the null hypothesis of no association. The red diagonal line indicates the expected null distribution. The histogram shows the distribution of raw *P* values across all tested variants, with the dashed horizontal line indicating the expected density under a uniform distribution.

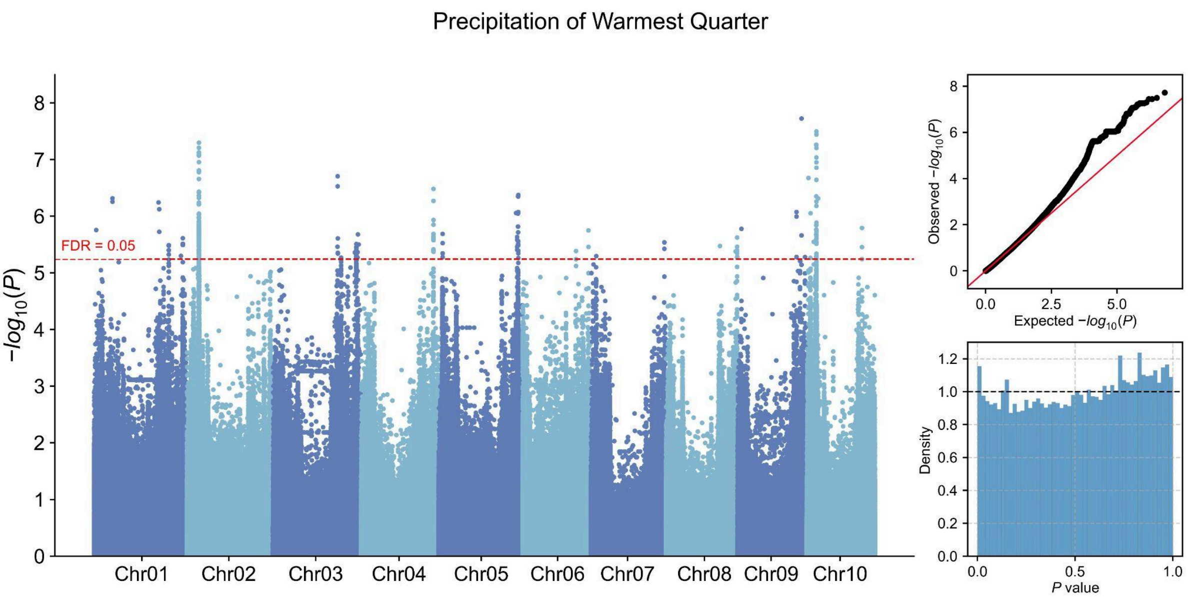

**Fig. S3. Genome-wide association results for precipitation of the warmest quarter.** The Manhattan plot shows variant associations across the genome, with each point representing a tested variant plotted by genomic position and −log10(*P*). Chromosomes are displayed in alternating colors, and the dashed red line denotes the false discovery rate (FDR) threshold of 0.05. The quantile–quantile plot compares observed and expected −log10(*P*) values under the null hypothesis of no association. The red diagonal line indicates the expected null distribution. The histogram shows the distribution of raw *P* values across all tested variants, with the dashed horizontal line indicating the expected density under a uniform distribution.

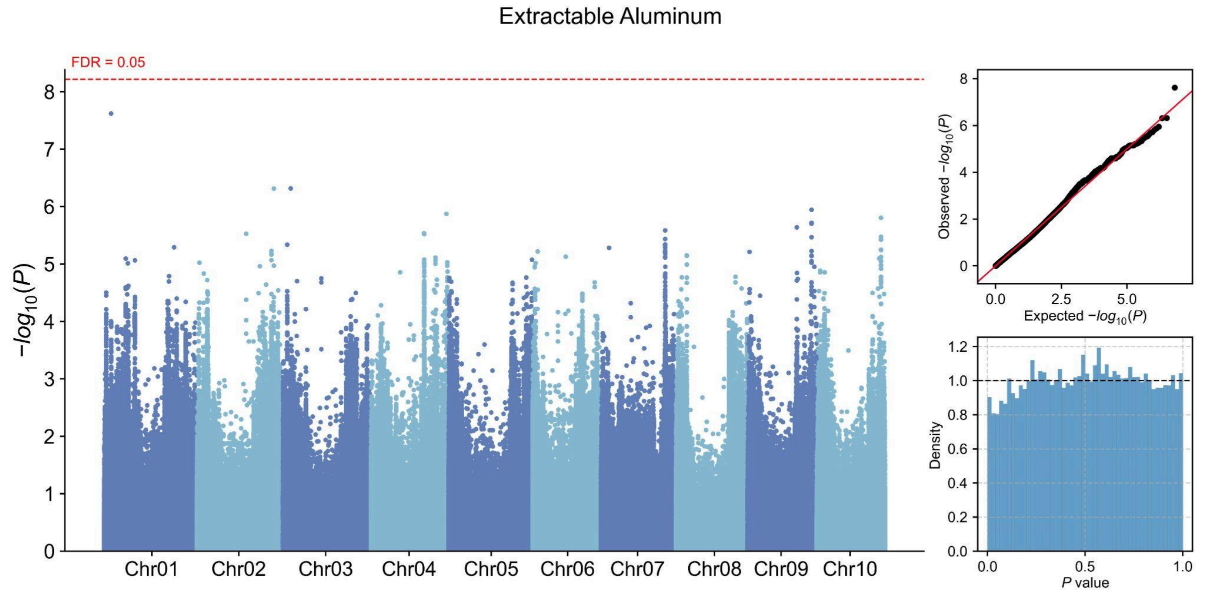

**Fig. S4. Genome-wide association results for soil extractable aluminum.** The Manhattan plot shows variant associations across the genome, with each point representing a tested variant plotted by genomic position and −log10($P$). Chromosomes are displayed in alternating colors, and the dashed red line denotes the false discovery rate (FDR) threshold of 0.05. The quantile–quantile plot compares observed and expected −log10($P$) values under the null hypothesis of no association. The red diagonal line indicates the expected null distribution. The histogram shows the distribution of raw $P$ values across all tested variants, with the dashed horizontal line indicating the expected density under a uniform distribution.

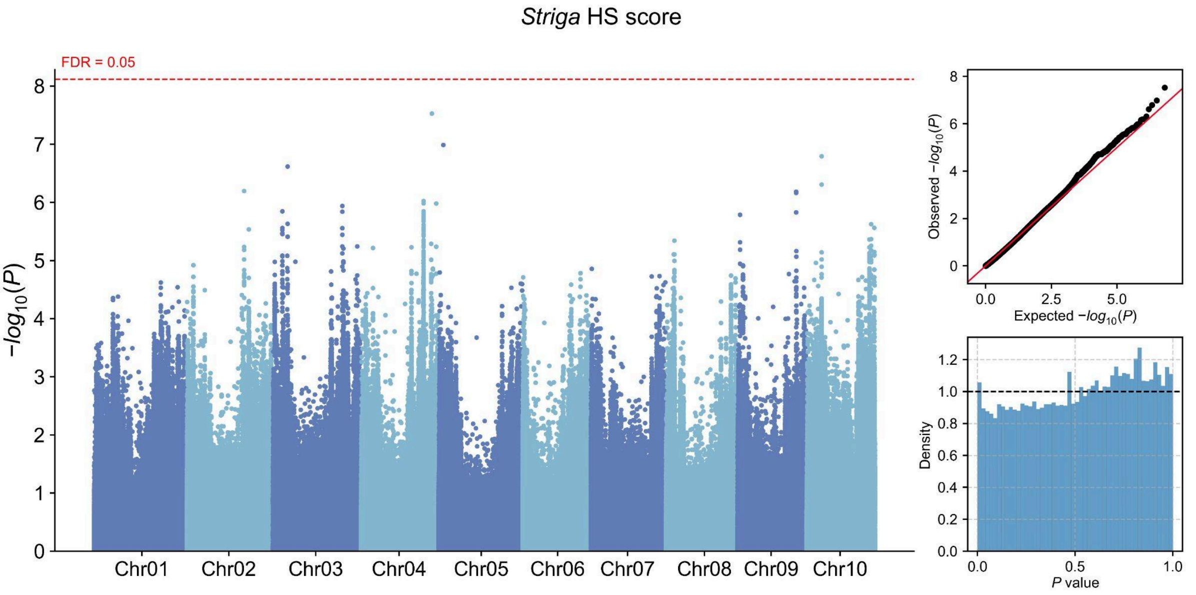

**Fig. S5. Genome-wide association results for *Striga* HS score.** The Manhattan plot shows variant associations across the genome, with each point representing a tested variant plotted by genomic position and −log10(*P*). Chromosomes are displayed in alternating colors, and the dashed red line denotes the false discovery rate (FDR) threshold of 0.05. The quantile–quantile plot compares observed and expected −log10(*P*) values under the null hypothesis of no association. The red diagonal line indicates the expected null distribution. The histogram shows the distribution of raw *P* values across all tested variants, with the dashed horizontal line indicating the expected density under a uniform distribution.

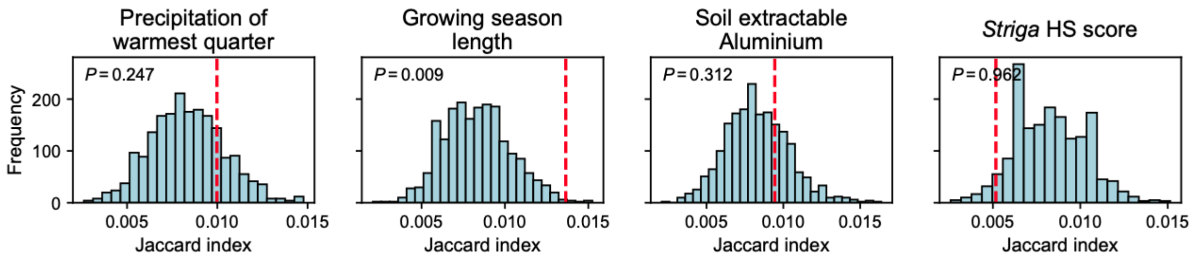

**Fig. S6. Enrichment of flowering-time genes among top 5% GEA genes.**
For each environmental variable, the red dashed line indicates the observed Jaccard index between flowering-time genes and the top 5% GEA candidate genes. Histograms show null distributions from 1,000 permutations of flowering-gene labels.

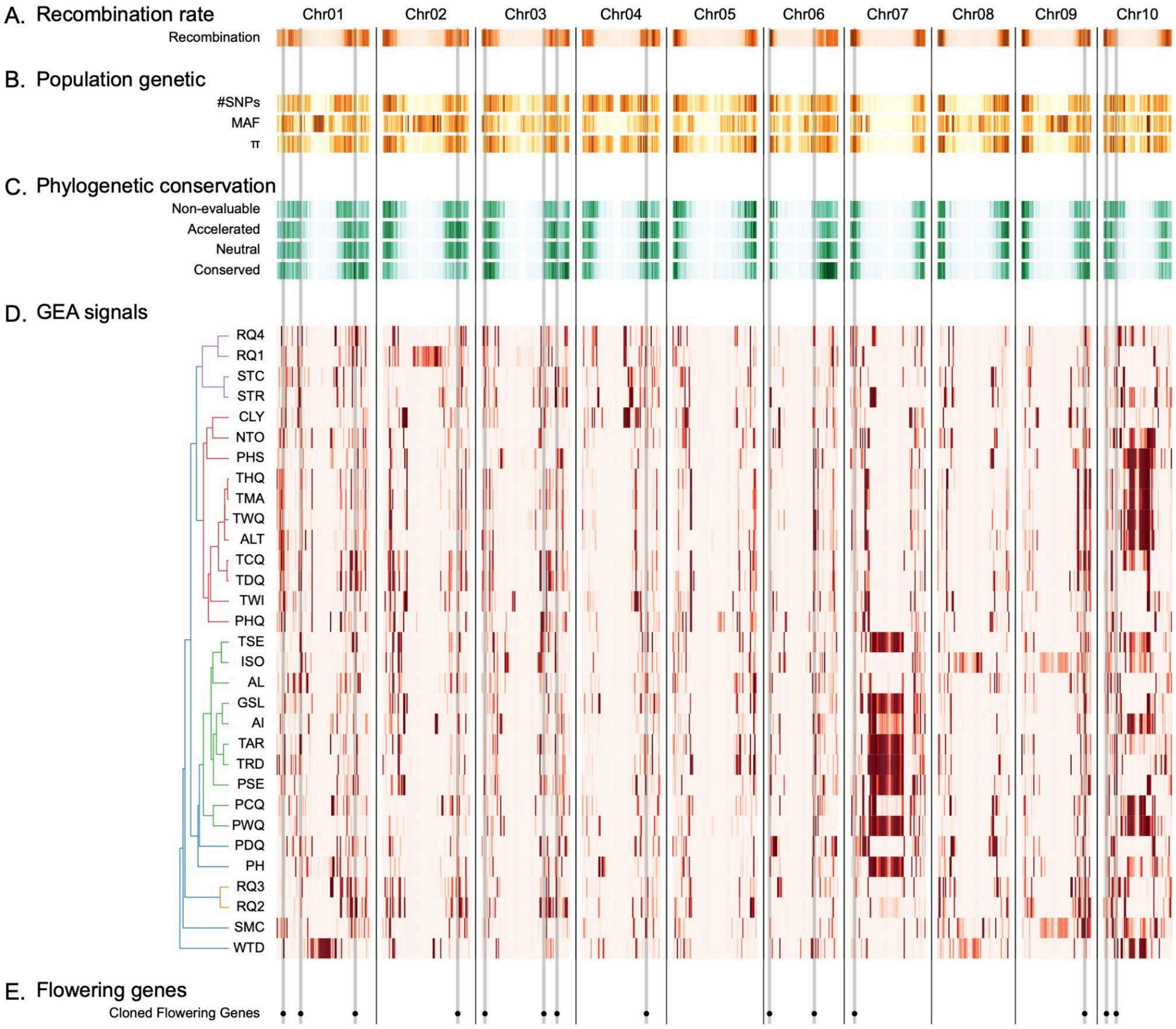

**Fig. S7. Additional genomic features in the diverse landscape of genomic features and genotype–environmental associations.** The ten sorghum chromosomes (Chr01–Chr10) were divided into equal-length bins. All tracks were *z*-scored within track; darker cells indicate higher-than-mean values, lighter cells indicate lower-than-mean values. (A) Recombination rate. (B) Population genetic features: SNP density, minor allele frequency (MAF), and nucleotide diversity ($\pi$). (C) Phylogenetic conservation categories: non-evaluable, accelerated, neutral, and conserved. (D) envGWAS: Each row corresponds to one environmental variable. The dendrogram clusters variables by similarity of their per-sample values across the resequenced landrace panel. The heat map shows enrichment of the top 5% significant SNPs across bins, calculated with a hypergeometric test using callable SNPs per bin as background, expressed as $-\log10(P)$, and *z*-scored within row. Darker cells indicate relative enrichment for that variable. Flowering-time genes are indicated along the x-axis; gray vertical bands span panels. (E) Genomic positions of cloned flowering-time genes. Gray vertical lines indicate the corresponding chromosome positions and extend through the upper genome-wide tracks.

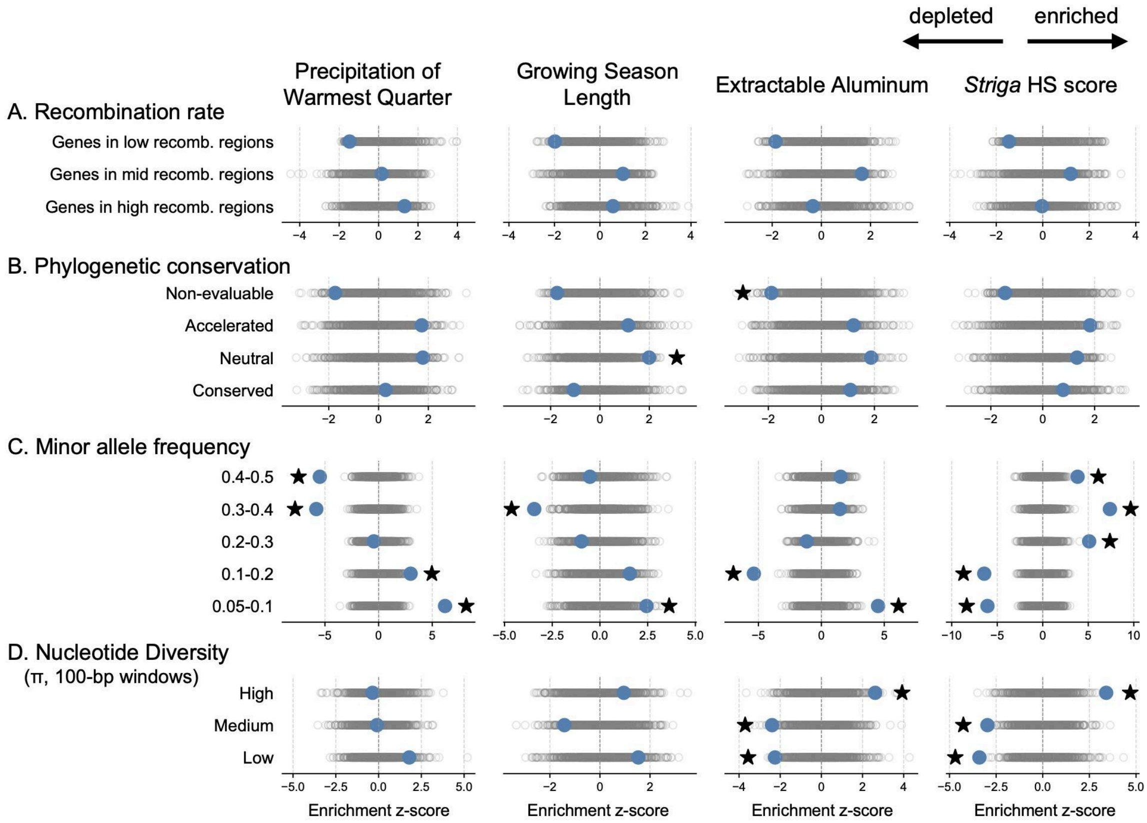

**Fig. S8. Additional enrichments of GEAs across categories of functional variation.** Enrichments were tested on the top 5% of GEAs for four environmental variables: precipitation of the warmest quarter, growing season length, extractable aluminum, and *Striga* HS score. Observed values (blue) are calculated as a *z*-score using 1,000 null permutations (gray). Significant enrichments are indicated by stars (two-sided empirical $P < 0.05$). (A) Recombination rate categories: genes in low, mid, and high recombination regions. (B) Phylogenetic conservation categories: non-evaluable, accelerated, neutral, and conserved sites. (C) MAF categories. (D) Nucleotide diversity ($\pi$) categories calculated in 100-kb windows.

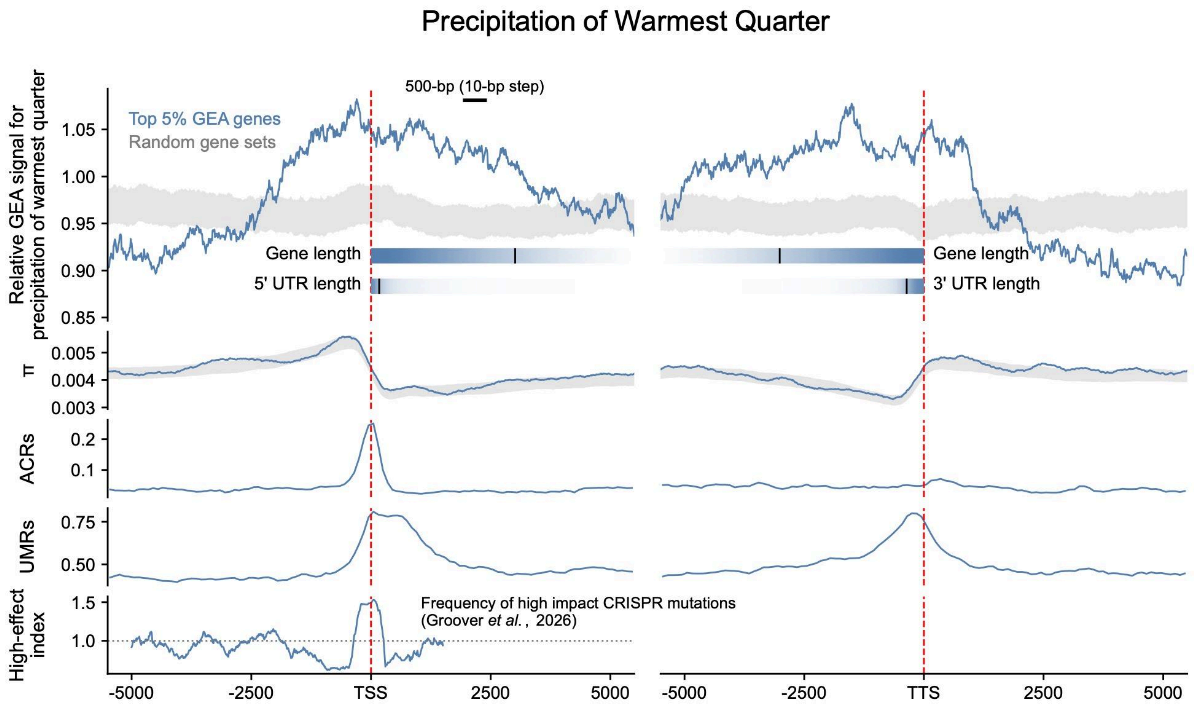

**Fig. S9. Spatial distribution of envGWAS signals around genes associated with precipitation of the warmest quarter.** The panel shows signal profiles around both ends of the top 5% of genes most strongly associated with precipitation of the warmest quarter. The x-axis indicates distance (bp) from the transcription start site (TSS; left) or transcription termination site (TTS; right). The bars labeled gene length and 5′/3′ UTR length show the distribution of gene and UTR lengths across genes, where darker shading indicates a higher proportion of genes extending to that position and black vertical ticks indicate the median length. The dark-blue curve shows the relative enrichment of envGWAS signals, calculated as the ratio of the first quartile (Q1) of −log10(*P*) values within sliding windows (500 bp window, 10 bp step) to the corresponding Q1 across the entire ±10 kb region around the TSS or TTS; the grey band denotes the 95% empirical range from 1,000 genome-wide random gene sets of equal size. The ACR and UMR tracks represent the proportion of bases overlapping each feature, and $\pi$ shows nucleotide diversity in the same windows. The bottom track shows a high-effect index, measuring the enrichment of high-effect CRISPR-accessible promoter mutations in each window from Groover et al. (2026) (dotted line, index=1: no enrichment).

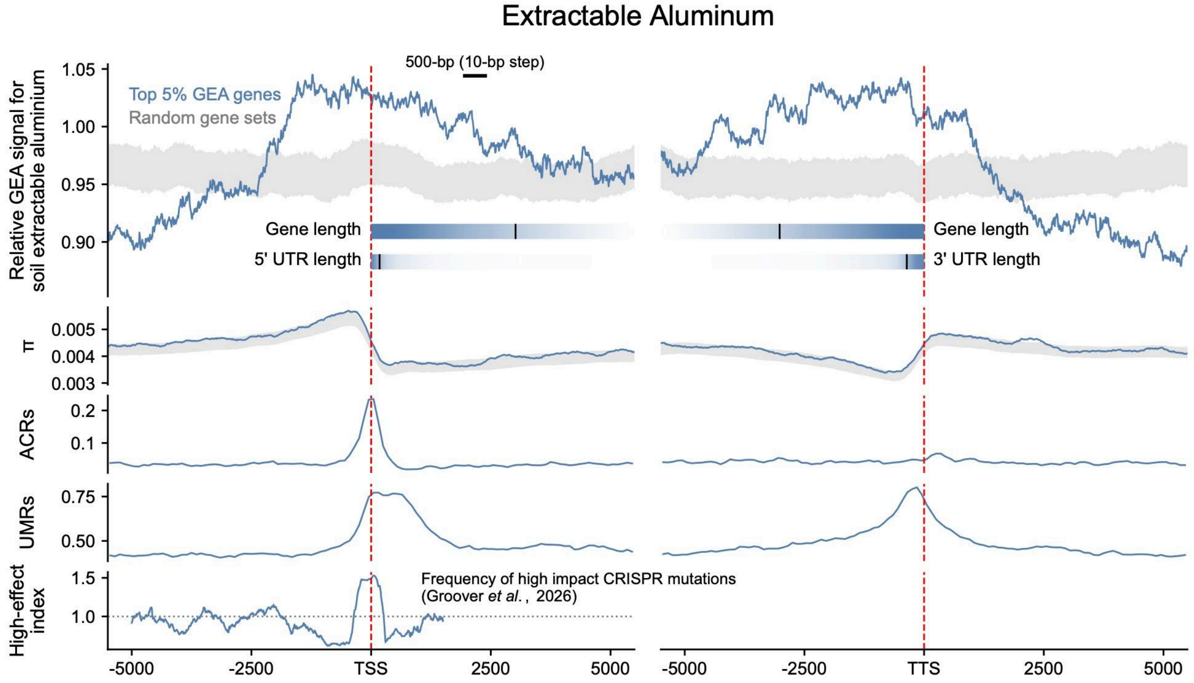

**Fig. S10. Spatial distribution of envGWAS signals around genes associated with soil extractable aluminum.** The panel shows signal profiles around both ends of the top 5% of genes most strongly associated with soil extractable aluminum. The x-axis indicates distance (bp) from the transcription start site (TSS; left) or transcription termination site (TTS; right). The bars labeled gene length and 5′/3′ UTR length show the distribution of gene and UTR lengths across genes, where darker shading indicates a higher proportion of genes extending to that position and black vertical ticks indicate the median length. The dark-blue curve shows the relative enrichment of envGWAS signals, calculated as the ratio of the first quartile (Q1) of −log10(*P*) values within sliding windows (500 bp window, 10 bp step) to the corresponding Q1 across the entire ±10 kb region around the TSS or TTS; the grey band denotes the 95% empirical range from 1,000 genome-wide random gene sets of equal size. The ACR and UMR tracks represent the proportion of bases overlapping each feature, and $\pi$ shows nucleotide diversity in the same windows. The bottom track shows a high-effect index, measuring the enrichment of high-effect CRISPR-accessible promoter mutations in each window from Groover et al. (2026) (dotted line, index=1: no enrichment).

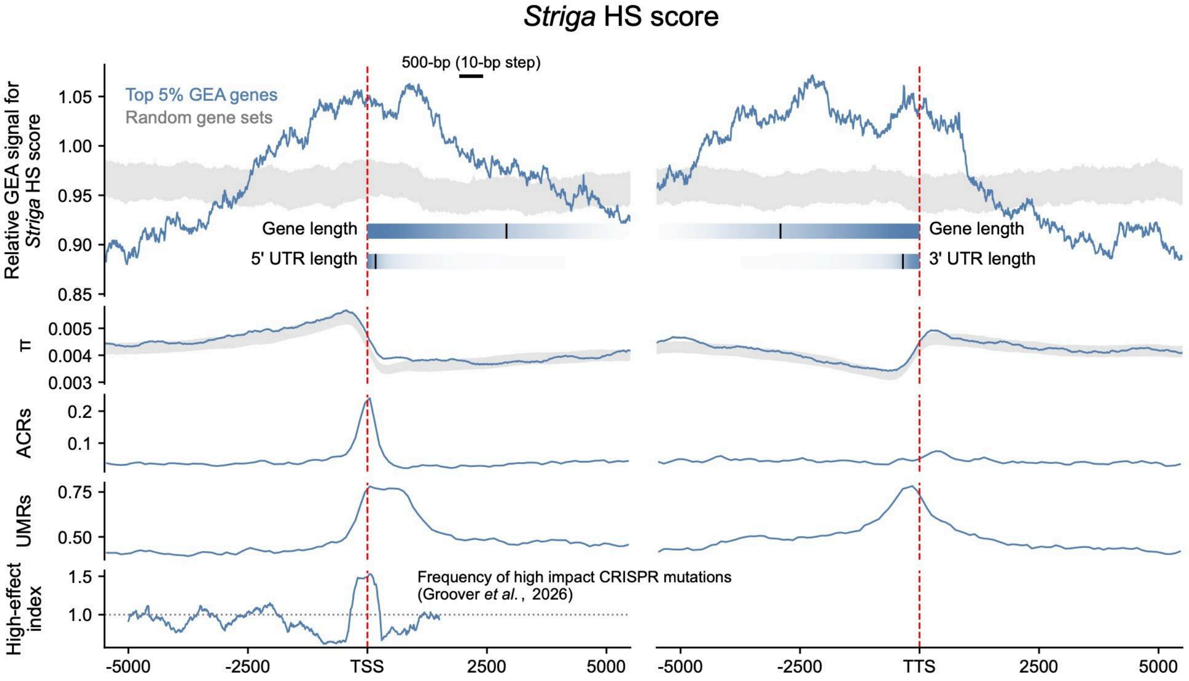

**Fig. S11. Spatial distribution of envGWAS signals around genes associated with *Striga* HS score.** The panel shows signal profiles around both ends of the top 5% of genes most strongly associated with *Striga* HS score. The x-axis indicates distance (bp) from the transcription start site (TSS; left) or transcription termination site (TTS; right). The bars labeled gene length and 5′/3′ UTR length show the distribution of gene and UTR lengths across genes, where darker shading indicates a higher proportion of genes extending to that position and black vertical ticks indicate the median length. The dark-blue curve shows the relative enrichment of envGWAS signals, calculated as the ratio of the first quartile (Q1) of −log10(*P*) values within sliding windows (500 bp window, 10 bp step) to the corresponding Q1 across the entire ±10 kb region around the TSS or TTS; the grey band denotes the 95% empirical range from 1,000 genome-wide random gene sets of equal size. The ACR and UMR tracks represent the proportion of bases overlapping each feature, and $\pi$ shows nucleotide diversity in the same windows. The bottom track shows a high-effect index, measuring the enrichment of high-effect CRISPR-accessible promoter mutations in each window from Groover et al. (2026) (dotted line, index=1: no enrichment).

### Tables

#### Table S1. Environmental variables for GEA

| Abbr. | Category | Variable Name | Unit | Data Source | Ref. |
|---|---|---|---|---|---|
| TMA | Temperature | Annual Mean Temperature | °C | CHELSA V2.1 | (5) |
| TRD | Temperature | Mean Diurnal Range | °C | CHELSA V2.1 | (5) |
| ISO | Temperature | Isothermality | % | CHELSA V2.1 | (5) |
| TSE | Temperature | Temperature Seasonality | °C | CHELSA V2.1 | (5) |
| TAR | Temperature | Temperature Annual Range | °C | CHELSA V2.1 | (5) |
| TWQ | Temperature | Mean Temperature of Wettest Quarter | °C | CHELSA V2.1 | (5) |
| TDQ | Temperature | Mean Temperature of Driest Quarter | °C | CHELSA V2.1 | (5) |
| THQ | Temperature | Mean Temperature of Warmest Quarter | °C | CHELSA V2.1 | (5) |
| TCQ | Temperature | Mean Temperature of Coldest Quarter | °C | CHELSA V2.1 | (5) |
| PSE | Precipitation | Precipitation Seasonality | mm | CHELSA V2.1 | (5) |
| PWQ | Precipitation | Precipitation of Wettest Quarter | mm | CHELSA V2.1 | (5) |
| PDQ | Precipitation | Precipitation of Driest Quarter | mm | CHELSA V2.1 | (5) |
| PHQ | Precipitation | Precipitation of Warmest Quarter | mm | CHELSA V2.1 | (5) |
| PCQ | Precipitation | Precipitation of Coldest Quarter | mm | CHELSA V2.1 | (5) |
| AI | Drought | Aridity Index | 1e−4 |  | (6) |
| GSL | Drought | Growing Season Length | days | CHELSA V2.1 | (5) |
| RQ1 | Radiation | Photosynthetically Active Radiation, January-March | W/m² | NASA |  |
| RQ2 | Radiation | Photosynthetically Active Radiation, April-June | W/m² | NASA |  |
| RQ3 | Radiation | Photosynthetically Active Radiation, July-September | W/m² | NASA |  |
| RQ4 | Radiation | Photosynthetically Active Radiation, October-December | W/m² | NASA |  |
| ALT | Topography | Altitude | m | WorldClim | (40) |
| TWI | Topography | Topographic wetness index |  | ENVIREM | (7) |
| AL | Soil & Hydrology | Extractable Aluminium | mg/kg | iSDAsoil | (41) |
| PH | Soil & Hydrology | Topsoil pH |  | iSDAsoil | (41) |
| NTO | Soil & Hydrology | Total nitrogen at 0-30 cm depth | g/kg | AfSoilGrids250m | (8) |
| PHS | Soil & Hydrology | Total phosphorus at 0-30 cm depth | mg/kg | AfSoilGrids250m | (8) |
| CLY | Soil & Hydrology | Clay fraction at 5 cm depth | % | SoilGrids250m | (9) |
| SMC | Soil & Hydrology | Extractable/Available Soil Moisture Capacity | cm | EarthData | (42) |
| WTD | Soil & Hydrology | Depth to Water Table | m |  | (10) |
| STR | *Striga* prevalence | *Striga* HS score |  | This study |  |

| | | | | |
|---|---|---|---|---|
| STC | *Striga* prevalence | *Striga* HS score (Crop) | | This study |

**Table S2. Site-level genomic contexts and enrichment statistics.**

Site-level enrichment was evaluated using site-level envGWAS results. For each environmental variable, sites were ranked by association *P* value and compared between each focal genomic context and the remaining background within the same analysis scope.

**Scope** defines the genomic background used for comparison, and **N sites** indicates the total number of sites within that scope. **Category** and **Context** define the genomic context being evaluated, and **N in context** indicates the number of sites assigned to that context. **Envir.** indicates the environmental variable tested.

Fold enrichment was quantified as the ratio of −log10(*P*) at a given threshold, top 5% or top 1%, inside versus outside the focal context. **Top5 obs.** and **Top1 obs.** report the observed enrichment values for the top 5% and top 1% most strongly associated sites, respectively. **Top5 null med.** / **Top1 null med.** and **Top5 null SD** / **Top1 null SD** summarize enrichment across 1,000 circular permutations of genomic context labels. **Top5 emp. P** / **Top1 emp. P** report empirical *P* values, and **Top5 BH FDR** / **Top1 BH FDR** report Benjamini–Hochberg adjusted *P* values. **Top5 Z** and **Top1 Z** are standardized enrichment scores relative to the null distribution; positive values indicate enrichment and negative values indicate depletion of GEA signals in the corresponding context.

Abbreviations: **PHQ**, precipitation of the warmest quarter; **GSL**, growing season length; **AL**, extractable aluminum; **STR**, *Striga* HS score.

| Scope | N sites | Category | Context | N in context | Envir. | Top5 obs. | Top5 null med. | Top5 null SD | Top5 emp *P* | Top5 BD FDR | Top5 *Z* | Top1 obs. | Top1 null med. | Top1 null SD | Top1 emp *P* | Top1 BD FDR | Top1 *Z* |
|---|---|---|---|---|---|---|---|---|---|---|---|---|---|---|---|---|---|
| Genome | 6,453,719 | Genomic | TEs | 3,125,648 | PHQ | 0.892 | 1.003 | 0.054 | 0.013 | 0.038 | -2.0 | 0.876 | 1.000 | 0.062 | 0.008 | 0.039 | -2.0 |
| Genome | 6,453,719 | Genomic | TEs | 3,125,648 | GSL | 0.885 | 0.999 | 0.061 | 0.017 | 0.044 | -1.9 | 0.899 | 0.999 | 0.055 | 0.014 | 0.056 | -1.9 |
| Genome | 6,453,719 | Genomic | TEs | 3,125,648 | AL | 0.836 | 1.001 | 0.064 | 0.003 | 0.016 | -2.6 | 0.854 | 0.999 | 0.053 | 0.005 | 0.038 | -2.8 |
| Genome | 6,453,719 | Genomic | TEs | 3,125,648 | STR | 0.889 | 1.001 | 0.048 | 0.004 | 0.018 | -2.3 | 0.912 | 0.999 | 0.043 | 0.027 | 0.068 | -2.1 |
| Genome | 6,453,719 | Genomic | Distal TEs | 2,578,954 | PHQ | 0.859 | 1.004 | 0.061 | 0.005 | 0.021 | -2.3 | 0.836 | 1.001 | 0.070 | 0.001 | 0.011 | -2.4 |
| Genome | 6,453,719 | Genomic | Distal TEs | 2,578,954 | GSL | 0.854 | 1.000 | 0.069 | 0.008 | 0.028 | -2.2 | 0.887 | 1.000 | 0.062 | 0.021 | 0.062 | -1.9 |

| | | | | | | | | | | | | | | | | | |
|---|---|---|---|---|---|---|---|---|---|---|---|---|---|---|---|---|---|
| Genome | 6,453,719 | Genomic | Distal TEs | 2,578,954 | AL | 0.801 | 1.003 | 0.073 | 0.003 | 0.016 | -2.8 | 0.809 | 1.000 | 0.060 | 0.003 | 0.024 | -3.2 |
| Genome | 6,453,719 | Genomic | Distal TEs | 2,578,954 | STR | 0.867 | 1.004 | 0.054 | 0.002 | 0.013 | -2.5 | 0.879 | 1.000 | 0.049 | 0.002 | 0.018 | -2.5 |
| Genome | 6,453,719 | Genomic | Prox. TEs | 546,694 | PHQ | 1.089 | 1.000 | 0.019 | 0.001 | 0.009 | 4.7 | 1.084 | 1.000 | 0.023 | 0.001 | 0.011 | 3.5 |
| Genome | 6,453,719 | Genomic | Prox. TEs | 546,694 | GSL | 1.056 | 0.998 | 0.021 | 0.012 | 0.037 | 2.6 | 1.029 | 1.000 | 0.021 | 0.072 | 0.132 | 1.4 |
| Genome | 6,453,719 | Genomic | Prox. TEs | 546,694 | AL | 1.086 | 0.998 | 0.022 | 0.006 | 0.023 | 3.8 | 1.077 | 1.000 | 0.021 | 0.006 | 0.038 | 3.5 |
| Genome | 6,453,719 | Genomic | Prox. TEs | 546,694 | STR | 1.077 | 0.998 | 0.019 | 0.001 | 0.009 | 3.9 | 1.083 | 0.999 | 0.019 | 0.001 | 0.011 | 4.4 |
| Genome | 6,453,719 | Genomic | Simp. reps. | 14,256 | PHQ | 1.051 | 1.001 | 0.022 | 0.013 | 0.038 | 2.2 | 1.009 | 1.001 | 0.032 | 0.403 | 0.428 | 0.2 |
| Genome | 6,453,719 | Genomic | Simp. reps. | 14,256 | GSL | 1.025 | 0.998 | 0.023 | 0.121 | 0.175 | 1.1 | 1.047 | 0.997 | 0.028 | 0.060 | 0.117 | 1.7 |
| Genome | 6,453,719 | Genomic | Simp. reps. | 14,256 | AL | 1.051 | 1.000 | 0.023 | 0.024 | 0.054 | 2.2 | 1.044 | 0.998 | 0.028 | 0.065 | 0.125 | 1.6 |
| Genome | 6,453,719 | Genomic | Simp. reps. | 14,256 | STR | 1.029 | 0.998 | 0.022 | 0.101 | 0.153 | 1.3 | 1.048 | 1.000 | 0.027 | 0.035 | 0.078 | 1.8 |
| Genome | 6,453,719 | Genomic | Other | 1,047,008 | PHQ | 1.018 | 0.999 | 0.014 | 0.087 | 0.138 | 1.4 | 1.027 | 0.999 | 0.018 | 0.092 | 0.160 | 1.4 |
| Genome | 6,453,719 | Genomic | Other | 1,047,008 | GSL | 1.007 | 0.997 | 0.015 | 0.299 | 0.345 | 0.5 | 1.036 | 1.000 | 0.015 | 0.007 | 0.038 | 2.5 |
| Genome | 6,453,719 | Genomic | Other | 1,047,008 | AL | 1.036 | 0.999 | 0.013 | 0.008 | 0.028 | 2.7 | 1.029 | 0.998 | 0.016 | 0.052 | 0.106 | 1.8 |
| Genome | 6,453,719 | Genomic | Other | 1,047,008 | STR | 1.012 | 1.000 | 0.015 | 0.191 | 0.238 | 0.8 | 1.029 | 0.999 | 0.015 | 0.037 | 0.081 | 1.9 |
| Genome | 6,453,719 | Genomic | Genic | 2,266,807 | PHQ | 1.114 | 0.998 | 0.056 | 0.014 | 0.039 | 2.0 | 1.121 | 0.999 | 0.065 | 0.016 | 0.057 | 1.9 |
| Genome | 6,453,719 | Genomic | Genic | 2,266,807 | GSL | 1.126 | 1.000 | 0.063 | 0.020 | 0.048 | 2.0 | 1.116 | 1.001 | 0.057 | 0.017 | 0.059 | 2.0 |
| Genome | 6,453,719 | Genomic | Genic | 2,266,807 | AL | 1.182 | 0.999 | 0.067 | 0.006 | 0.023 | 2.7 | 1.143 | 1.000 | 0.055 | 0.007 | 0.038 | 2.6 |
| Genome | 6,453,719 | Genomic | Genic | 2,266,807 | STR | 1.119 | 1.000 | 0.050 | 0.017 | 0.044 | 2.4 | 1.083 | 1.001 | 0.045 | 0.047 | 0.097 | 1.9 |
| Genome | 6,453,719 | Epigen. | ACRs | 104,250 | PHQ | 1.114 | 1.003 | 0.047 | 0.001 | 0.009 | 2.5 | 1.094 | 1.002 | 0.056 | 0.023 | 0.063 | 1.7 |

| | | | | | | | | | | | | | | | | | |
|---|---|---|---|---|---|---|---|---|---|---|---|---|---|---|---|---|---|
| Genome | 6,453,719 | Epigen. | ACRs | 104,250 | GSL | 1.090 | 1.002 | 0.056 | 0.037 | 0.075 | 1.6 | 1.069 | 1.001 | 0.051 | 0.059 | 0.116 | 1.4 |
| Genome | 6,453,719 | Epigen. | ACRs | 104,250 | AL | 1.127 | 1.005 | 0.057 | 0.003 | 0.016 | 2.2 | 1.096 | 1.002 | 0.051 | 0.016 | 0.057 | 1.9 |
| Genome | 6,453,719 | Epigen. | ACRs | 104,250 | STR | 1.112 | 1.004 | 0.044 | 0.003 | 0.016 | 2.5 | 1.087 | 1.001 | 0.041 | 0.008 | 0.039 | 2.2 |
| Genome | 6,453,719 | Epigen. | Genic ACRs | 79,998 | PHQ | 1.128 | 1.002 | 0.048 | 0.001 | 0.009 | 2.7 | 1.102 | 1.001 | 0.057 | 0.023 | 0.063 | 1.8 |
| Genome | 6,453,719 | Epigen. | Genic ACRs | 79,998 | GSL | 1.076 | 1.002 | 0.058 | 0.089 | 0.138 | 1.3 | 1.053 | 1.002 | 0.053 | 0.150 | 0.246 | 1.0 |
| Genome | 6,453,719 | Epigen. | Genic ACRs | 79,998 | AL | 1.123 | 1.005 | 0.059 | 0.007 | 0.026 | 2.1 | 1.085 | 1.002 | 0.053 | 0.025 | 0.064 | 1.7 |
| Genome | 6,453,719 | Epigen. | Genic ACRs | 79,998 | STR | 1.091 | 1.005 | 0.045 | 0.015 | 0.040 | 2.0 | 1.071 | 1.000 | 0.043 | 0.045 | 0.094 | 1.7 |
| Genome | 6,453,719 | Epigen. | Distal ACRs | 24,252 | PHQ | 1.047 | 0.999 | 0.048 | 0.177 | 0.227 | 1.0 | 1.064 | 0.999 | 0.065 | 0.173 | 0.264 | 1.0 |
| Genome | 6,453,719 | Epigen. | Distal ACRs | 24,252 | GSL | 1.116 | 0.999 | 0.057 | 0.023 | 0.052 | 2.0 | 1.099 | 0.997 | 0.057 | 0.035 | 0.078 | 1.8 |
| Genome | 6,453,719 | Epigen. | Distal ACRs | 24,252 | AL | 1.137 | 1.001 | 0.055 | 0.004 | 0.018 | 2.5 | 1.123 | 0.998 | 0.060 | 0.021 | 0.062 | 2.1 |
| Genome | 6,453,719 | Epigen. | Distal ACRs | 24,252 | STR | 1.167 | 0.997 | 0.047 | 0.002 | 0.013 | 3.5 | 1.122 | 0.997 | 0.049 | 0.006 | 0.038 | 2.6 |
| Genome | 6,453,719 | Epigen. | UMRs | 1,311,928 | PHQ | 1.098 | 0.998 | 0.047 | 0.009 | 0.030 | 2.1 | 1.093 | 1.000 | 0.055 | 0.021 | 0.062 | 1.7 |
| Genome | 6,453,719 | Epigen. | UMRs | 1,311,928 | GSL | 1.095 | 1.001 | 0.054 | 0.040 | 0.078 | 1.8 | 1.096 | 0.999 | 0.049 | 0.015 | 0.057 | 2.0 |
| Genome | 6,453,719 | Epigen. | UMRs | 1,311,928 | AL | 1.142 | 0.999 | 0.057 | 0.004 | 0.018 | 2.5 | 1.100 | 0.999 | 0.047 | 0.019 | 0.062 | 2.2 |
| Genome | 6,453,719 | Epigen. | UMRs | 1,311,928 | STR | 1.107 | 1.002 | 0.043 | 0.005 | 0.021 | 2.5 | 1.080 | 1.001 | 0.038 | 0.025 | 0.064 | 2.1 |
| Genome | 6,453,719 | Epigen. | Genic UMRs | 954,838 | PHQ | 1.086 | 0.999 | 0.049 | 0.018 | 0.045 | 1.8 | 1.087 | 1.000 | 0.057 | 0.033 | 0.076 | 1.6 |
| Genome | 6,453,719 | Epigen. | Genic UMRs | 954,838 | GSL | 1.088 | 1.001 | 0.056 | 0.062 | 0.108 | 1.6 | 1.082 | 1.000 | 0.051 | 0.023 | 0.063 | 1.7 |
| Genome | 6,453,719 | Epigen. | Genic UMRs | 954,838 | AL | 1.144 | 1.000 | 0.059 | 0.004 | 0.018 | 2.5 | 1.097 | 1.001 | 0.050 | 0.015 | 0.057 | 2.0 |
| Genome | 6,453,719 | Epigen. | Genic UMRs | 954,838 | STR | 1.105 | 1.003 | 0.044 | 0.006 | 0.023 | 2.4 | 1.073 | 1.002 | 0.039 | 0.033 | 0.076 | 1.9 |

| | | | | | | | | | | | | | | | | | |
|---|---|---|---|---|---|---|---|---|---|---|---|---|---|---|---|---|---|
| Genome | 6,453,719 | Epigen. | Distal UMRs | 357,090 | PHQ | 1.090 | 1.001 | 0.032 | 0.003 | 0.016 | 2.9 | 1.074 | 1.000 | 0.038 | 0.024 | 0.064 | 2.0 |
| Genome | 6,453,719 | Epigen. | Distal UMRs | 357,090 | GSL | 1.075 | 0.999 | 0.036 | 0.020 | 0.048 | 2.1 | 1.104 | 0.999 | 0.034 | 0.002 | 0.018 | 3.1 |
| Genome | 6,453,719 | Epigen. | Distal UMRs | 357,090 | AL | 1.101 | 0.999 | 0.038 | 0.014 | 0.039 | 2.7 | 1.071 | 0.998 | 0.034 | 0.023 | 0.063 | 2.1 |
| Genome | 6,453,719 | Epigen. | Distal UMRs | 357,090 | STR | 1.073 | 0.997 | 0.030 | 0.015 | 0.040 | 2.4 | 1.072 | 0.999 | 0.029 | 0.007 | 0.038 | 2.6 |
| CDS | 241,038 | CDS | Missense | 112,470 | PHQ | 0.989 | 1.000 | 0.009 | 0.103 | 0.154 | -1.3 | 0.996 | 1.001 | 0.013 | 0.329 | 0.386 | -0.4 |
| CDS | 241,038 | CDS | Missense | 112,470 | GSL | 0.998 | 1.000 | 0.007 | 0.401 | 0.417 | -0.2 | 1.009 | 1.000 | 0.014 | 0.237 | 0.327 | 0.7 |
| CDS | 241,038 | CDS | Missense | 112,470 | AL | 0.997 | 1.000 | 0.008 | 0.350 | 0.384 | -0.4 | 1.007 | 1.000 | 0.012 | 0.267 | 0.339 | 0.6 |
| CDS | 241,038 | CDS | Missense | 112,470 | STR | 0.989 | 1.000 | 0.008 | 0.089 | 0.138 | -1.4 | 1.007 | 0.999 | 0.012 | 0.274 | 0.339 | 0.6 |
| CDS | 241,038 | CDS | Syn. | 113,948 | PHQ | 1.009 | 1.000 | 0.009 | 0.161 | 0.216 | 1.0 | 1.003 | 0.999 | 0.013 | 0.378 | 0.410 | 0.3 |
| CDS | 241,038 | CDS | Syn. | 113,948 | GSL | 1.006 | 1.000 | 0.008 | 0.211 | 0.259 | 0.8 | 0.991 | 1.000 | 0.014 | 0.268 | 0.339 | -0.6 |
| CDS | 241,038 | CDS | Syn. | 113,948 | AL | 1.009 | 0.999 | 0.009 | 0.164 | 0.217 | 1.0 | 0.992 | 0.999 | 0.012 | 0.248 | 0.331 | -0.6 |
| CDS | 241,038 | CDS | Syn. | 113,948 | STR | 1.010 | 1.000 | 0.008 | 0.116 | 0.170 | 1.2 | 0.991 | 1.001 | 0.012 | 0.224 | 0.317 | -0.8 |
| CDS | 241,038 | CDS | LoF | 6,968 | PHQ | 1.021 | 1.000 | 0.022 | 0.152 | 0.207 | 1.0 | 1.003 | 0.997 | 0.034 | 0.437 | 0.454 | 0.1 |
| CDS | 241,038 | CDS | LoF | 6,968 | GSL | 0.968 | 0.998 | 0.022 | 0.065 | 0.111 | -1.5 | 0.961 | 0.996 | 0.036 | 0.167 | 0.263 | -1.0 |
| CDS | 241,038 | CDS | LoF | 6,968 | AL | 0.961 | 1.001 | 0.023 | 0.032 | 0.067 | -1.8 | 1.009 | 1.000 | 0.034 | 0.380 | 0.410 | 0.3 |
| CDS | 241,038 | CDS | LoF | 6,968 | STR | 1.005 | 0.999 | 0.023 | 0.392 | 0.417 | 0.3 | 1.018 | 0.992 | 0.029 | 0.221 | 0.316 | 0.8 |
| Genic (±15 kb) | 3,883,479 | Pop. gen. | MAF 0.05-0.10 | 1,024,835 | PHQ | 1.055 | 1.000 | 0.009 | 0.001 | 0.009 | 6.2 | 1.111 | 0.999 | 0.011 | 0.001 | 0.011 | 9.3 |
| Genic (±15 kb) | 3,883,479 | Pop. gen. | MAF 0.05-0.10 | 1,024,835 | GSL | 1.020 | 1.000 | 0.008 | 0.012 | 0.037 | 2.4 | 1.005 | 1.001 | 0.011 | 0.343 | 0.392 | 0.4 |

| | | | | | | | | | | | | | | | | | |
|---|---|---|---|---|---|---|---|---|---|---|---|---|---|---|---|---|---|
| Genic (±15 kb) | 3,883,479 | Pop. gen. | MAF 0.05-0.10 | 1,024,835 | AL | 1.040 | 1.000 | 0.009 | 0.001 | 0.009 | 4.5 | 1.094 | 1.000 | 0.012 | 0.001 | 0.011 | 7.4 |
| Genic (±15 kb) | 3,883,479 | Pop. gen. | MAF 0.05-0.10 | 1,024,835 | STR | 0.949 | 1.000 | 0.008 | 0.001 | 0.009 | -6.1 | 0.980 | 1.000 | 0.010 | 0.030 | 0.073 | -1.9 |
| Genic (±15 kb) | 3,883,479 | Pop. gen. | MAF 0.10-0.20 | 1,182,414 | PHQ | 1.024 | 1.000 | 0.008 | 0.002 | 0.013 | 3.0 | 1.029 | 1.000 | 0.011 | 0.008 | 0.039 | 2.6 |
| Genic (±15 kb) | 3,883,479 | Pop. gen. | MAF 0.10-0.20 | 1,182,414 | GSL | 1.012 | 1.000 | 0.007 | 0.061 | 0.108 | 1.6 | 1.035 | 0.999 | 0.010 | 0.001 | 0.011 | 3.4 |
| Genic (±15 kb) | 3,883,479 | Pop. gen. | MAF 0.10-0.20 | 1,182,414 | AL | 0.956 | 1.000 | 0.008 | 0.001 | 0.009 | -5.4 | 0.947 | 0.999 | 0.012 | 0.001 | 0.011 | -4.4 |
| Genic (±15 kb) | 3,883,479 | Pop. gen. | MAF 0.10-0.20 | 1,182,414 | STR | 0.948 | 1.000 | 0.008 | 0.001 | 0.009 | -6.4 | 0.960 | 1.000 | 0.010 | 0.001 | 0.011 | -4.0 |
| Genic (±15 kb) | 3,883,479 | Pop. gen. | MAF 0.20-0.30 | 657,107 | PHQ | 0.996 | 1.000 | 0.009 | 0.335 | 0.370 | -0.4 | 0.971 | 1.000 | 0.013 | 0.011 | 0.048 | -2.3 |
| Genic (±15 kb) | 3,883,479 | Pop. gen. | MAF 0.20-0.30 | 657,107 | GSL | 0.992 | 1.000 | 0.009 | 0.162 | 0.216 | -1.0 | 0.971 | 1.000 | 0.012 | 0.007 | 0.038 | -2.4 |
| Genic (±15 kb) | 3,883,479 | Pop. gen. | MAF 0.20-0.30 | 657,107 | AL | 0.988 | 0.999 | 0.010 | 0.124 | 0.176 | -1.2 | 0.958 | 0.999 | 0.014 | 0.001 | 0.011 | -3.0 |
| Genic (±15 kb) | 3,883,479 | Pop. gen. | MAF 0.20-0.30 | 657,107 | STR | 1.048 | 1.000 | 0.009 | 0.001 | 0.009 | 5.1 | 1.048 | 1.000 | 0.012 | 0.001 | 0.011 | 4.1 |
| Genic (±15 kb) | 3,883,479 | Pop. gen. | MAF 0.30-0.40 | 532,776 | PHQ | 0.939 | 1.000 | 0.010 | 0.001 | 0.009 | -5.8 | 0.894 | 0.999 | 0.015 | 0.001 | 0.011 | -7.1 |
| Genic (±15 kb) | 3,883,479 | Pop. gen. | MAF 0.30-0.40 | 532,776 | GSL | 0.967 | 1.000 | 0.010 | 0.002 | 0.013 | -3.4 | 0.974 | 1.000 | 0.013 | 0.031 | 0.074 | -2.0 |
| Genic (±15 kb) | 3,883,479 | Pop. gen. | MAF 0.30-0.40 | 532,776 | AL | 1.016 | 1.000 | 0.011 | 0.072 | 0.118 | 1.5 | 0.995 | 0.999 | 0.016 | 0.379 | 0.410 | -0.4 |

| | | | | | | | | | | | | | | | | | |
|---|---|---|---|---|---|---|---|---|---|---|---|---|---|---|---|---|---|
| Genic (±15 kb) | 3,883,479 | Pop. gen. | MAF 0.30-0.40 | 532,776 | STR | 1.083 | 1.000 | 0.011 | 0.001 | 0.009 | 7.4 | 1.034 | 1.000 | 0.014 | 0.010 | 0.045 | 2.5 |
| Genic (±15 kb) | 3,883,479 | Pop. gen. | MAF 0.40-0.50 | 486,347 | PHQ | 0.934 | 1.000 | 0.012 | 0.001 | 0.009 | -5.5 | 0.894 | 0.999 | 0.017 | 0.001 | 0.011 | -6.1 |
| Genic (±15 kb) | 3,883,479 | Pop. gen. | MAF 0.40-0.50 | 486,347 | GSL | 0.994 | 1.000 | 0.011 | 0.306 | 0.346 | -0.5 | 0.980 | 0.999 | 0.016 | 0.094 | 0.162 | -1.2 |
| Genic (±15 kb) | 3,883,479 | Pop. gen. | MAF 0.40-0.50 | 486,347 | AL | 1.018 | 1.000 | 0.012 | 0.068 | 0.114 | 1.5 | 1.006 | 0.999 | 0.018 | 0.357 | 0.401 | 0.3 |
| Genic (±15 kb) | 3,883,479 | Pop. gen. | MAF 0.40-0.50 | 486,347 | STR | 1.046 | 1.000 | 0.012 | 0.001 | 0.009 | 3.8 | 0.998 | 1.000 | 0.014 | 0.459 | 0.466 | -0.1 |
| Genic (±15 kb) | 3,883,479 | Pop. gen. | Low π | 209,488 | PHQ | 1.022 | 0.999 | 0.013 | 0.048 | 0.089 | 1.8 | 1.019 | 0.999 | 0.017 | 0.149 | 0.246 | 1.0 |
| Genic (±15 kb) | 3,883,479 | Pop. gen. | Low π | 209,488 | GSL | 1.019 | 1.000 | 0.012 | 0.070 | 0.116 | 1.5 | 1.001 | 0.999 | 0.016 | 0.457 | 0.466 | 0.1 |
| Genic (±15 kb) | 3,883,479 | Pop. gen. | Low π | 209,488 | AL | 0.970 | 1.000 | 0.013 | 0.013 | 0.038 | -2.3 | 0.985 | 1.000 | 0.017 | 0.181 | 0.271 | -0.9 |
| Genic (±15 kb) | 3,883,479 | Pop. gen. | Low π | 209,488 | STR | 0.955 | 1.000 | 0.013 | 0.001 | 0.009 | -3.4 | 0.989 | 0.999 | 0.015 | 0.230 | 0.322 | -0.7 |
| Genic (±15 kb) | 3,883,479 | Pop. gen. | Med. π | 1,420,115 | PHQ | 0.999 | 1.000 | 0.010 | 0.450 | 0.453 | -0.1 | 0.998 | 1.001 | 0.013 | 0.407 | 0.429 | -0.2 |
| Genic (±15 kb) | 3,883,479 | Pop. gen. | Med. π | 1,420,115 | GSL | 0.987 | 1.000 | 0.009 | 0.064 | 0.110 | -1.4 | 0.980 | 1.001 | 0.012 | 0.040 | 0.085 | -1.7 |
| Genic (±15 kb) | 3,883,479 | Pop. gen. | Med. π | 1,420,115 | AL | 0.974 | 1.000 | 0.011 | 0.012 | 0.037 | -2.4 | 0.987 | 1.000 | 0.014 | 0.164 | 0.262 | -1.0 |
| Genic (±15 kb) | 3,883,479 | Pop. gen. | Med. π | 1,420,115 | STR | 0.972 | 1.000 | 0.010 | 0.002 | 0.013 | -3.0 | 0.988 | 1.000 | 0.011 | 0.158 | 0.256 | -1.0 |

| | | | | | | | | | | | | | | | | | |
|---|---|---|---|---|---|---|---|---|---|---|---|---|---|---|---|---|---|
| Genic (±15 kb) | 3,883,479 | Pop. gen. | High π | 2,253,876 | PHQ | 0.996 | 1.000 | 0.011 | 0.378 | 0.408 | -0.3 | 0.997 | 1.000 | 0.014 | 0.466 | 0.469 | -0.1 |
| Genic (±15 kb) | 3,883,479 | Pop. gen. | High π | 2,253,876 | GSL | 1.009 | 1.000 | 0.010 | 0.190 | 0.238 | 1.0 | 1.020 | 1.000 | 0.014 | 0.068 | 0.128 | 1.5 |
| Genic (±15 kb) | 3,883,479 | Pop. gen. | High π | 2,253,876 | AL | 1.032 | 1.000 | 0.012 | 0.009 | 0.030 | 2.6 | 1.015 | 1.000 | 0.016 | 0.168 | 0.263 | 1.0 |
| Genic (±15 kb) | 3,883,479 | Pop. gen. | High π | 2,253,876 | STR | 1.038 | 1.000 | 0.011 | 0.001 | 0.009 | 3.4 | 1.014 | 1.000 | 0.013 | 0.150 | 0.246 | 1.1 |
| Genic (±15 kb) | 3,883,479 | Phylo. | Conserved | 80,068 | PHQ | 1.003 | 1.000 | 0.010 | 0.387 | 0.414 | 0.3 | 0.994 | 1.000 | 0.014 | 0.287 | 0.349 | -0.5 |
| Genic (±15 kb) | 3,883,479 | Phylo. | Conserved | 80,068 | GSL | 0.990 | 1.000 | 0.010 | 0.138 | 0.192 | -1.1 | 0.993 | 1.000 | 0.014 | 0.321 | 0.380 | -0.5 |
| Genic (±15 kb) | 3,883,479 | Phylo. | Conserved | 80,068 | AL | 1.010 | 1.000 | 0.009 | 0.141 | 0.194 | 1.1 | 1.008 | 1.000 | 0.013 | 0.271 | 0.339 | 0.6 |
| Genic (±15 kb) | 3,883,479 | Phylo. | Conserved | 80,068 | STR | 1.007 | 0.999 | 0.010 | 0.215 | 0.261 | 0.8 | 1.017 | 1.000 | 0.013 | 0.089 | 0.157 | 1.4 |
| Genic (±15 kb) | 3,883,479 | Phylo. | Accelerated | 182,730 | PHQ | 1.013 | 1.000 | 0.007 | 0.043 | 0.082 | 1.7 | 1.015 | 1.000 | 0.009 | 0.071 | 0.132 | 1.5 |
| Genic (±15 kb) | 3,883,479 | Phylo. | Accelerated | 182,730 | GSL | 1.008 | 1.000 | 0.007 | 0.129 | 0.181 | 1.1 | 1.018 | 1.000 | 0.010 | 0.019 | 0.062 | 1.9 |
| Genic (±15 kb) | 3,883,479 | Phylo. | Accelerated | 182,730 | AL | 1.008 | 1.000 | 0.007 | 0.095 | 0.145 | 1.2 | 1.006 | 1.000 | 0.010 | 0.268 | 0.339 | 0.6 |
| Genic (±15 kb) | 3,883,479 | Phylo. | Accelerated | 182,730 | STR | 1.011 | 1.000 | 0.006 | 0.027 | 0.058 | 1.8 | 0.993 | 1.000 | 0.008 | 0.173 | 0.264 | -0.9 |
| Genic (±15 kb) | 3,883,479 | Phylo. | Neutral | 1,615,237 | PHQ | 1.016 | 1.000 | 0.009 | 0.038 | 0.076 | 1.8 | 1.023 | 1.001 | 0.010 | 0.010 | 0.045 | 2.4 |

| | | | | | | | | | | | | | | | | | |
|---|---|---|---|---|---|---|---|---|---|---|---|---|---|---|---|---|---|
| Genic (±15 kb) | 3,883,479 | Phylo. | Neutral | 1,615,237 | GSL | 1.015 | 1.000 | 0.008 | 0.019 | 0.047 | 2.0 | 1.021 | 1.000 | 0.011 | 0.018 | 0.061 | 1.9 |
| Genic (±15 kb) | 3,883,479 | Phylo. | Neutral | 1,615,237 | AL | 1.015 | 1.000 | 0.008 | 0.029 | 0.062 | 1.9 | 1.010 | 1.000 | 0.010 | 0.196 | 0.287 | 0.9 |
| Genic (±15 kb) | 3,883,479 | Phylo. | Neutral | 1,615,237 | STR | 1.009 | 1.001 | 0.007 | 0.082 | 0.131 | 1.3 | 1.005 | 1.000 | 0.008 | 0.272 | 0.339 | 0.6 |
| Genic (±15 kb) | 3,883,479 | Phylo. | Non-evaluable | 2,005,444 | PHQ | 0.982 | 1.000 | 0.010 | 0.039 | 0.077 | -1.7 | 0.976 | 0.999 | 0.011 | 0.014 | 0.056 | -2.1 |
| Genic (±15 kb) | 3,883,479 | Phylo. | Non-evaluable | 2,005,444 | GSL | 0.985 | 0.999 | 0.009 | 0.037 | 0.075 | -1.7 | 0.977 | 1.000 | 0.013 | 0.020 | 0.062 | -1.8 |
| Genic (±15 kb) | 3,883,479 | Phylo. | Non-evaluable | 2,005,444 | AL | 0.984 | 0.999 | 0.009 | 0.025 | 0.055 | -1.9 | 0.989 | 0.999 | 0.012 | 0.186 | 0.275 | -0.9 |
| Genic (±15 kb) | 3,883,479 | Phylo. | Non-evaluable | 2,005,444 | STR | 0.989 | 0.999 | 0.007 | 0.062 | 0.108 | -1.5 | 0.995 | 1.000 | 0.009 | 0.310 | 0.370 | -0.5 |
| Gene Body | 858,402 | RiboSNitch | RiboSNitch | 240,854 | PHQ | 1.001 | 1.000 | 0.004 | 0.420 | 0.426 | 0.2 | 0.995 | 1.000 | 0.005 | 0.256 | 0.335 | -0.9 |
| Gene Body | 858,402 | RiboSNitch | RiboSNitch | 240,854 | GSL | 0.997 | 1.000 | 0.003 | 0.173 | 0.224 | -1.0 | 0.987 | 0.999 | 0.006 | 0.014 | 0.056 | -2.2 |
| Gene Body | 858,402 | RiboSNitch | RiboSNitch | 240,854 | AL | 0.997 | 1.000 | 0.004 | 0.173 | 0.224 | -0.9 | 0.997 | 1.000 | 0.006 | 0.346 | 0.392 | -0.4 |
| Gene Body | 858,402 | RiboSNitch | RiboSNitch | 240,854 | STR | 0.997 | 1.000 | 0.004 | 0.255 | 0.302 | -0.7 | 1.007 | 1.000 | 0.005 | 0.055 | 0.110 | 1.5 |
| Genic (±2 kb) | 2,266,807 | Gene struct. | Upstream (2K) | 831,210 | PHQ | 1.002 | 1.000 | 0.007 | 0.407 | 0.417 | 0.2 | 0.993 | 1.001 | 0.010 | 0.255 | 0.335 | -0.8 |
| Genic (±2 kb) | 2,266,807 | Gene struct. | Upstream (2K) | 831,210 | GSL | 1.009 | 1.000 | 0.007 | 0.111 | 0.164 | 1.2 | 1.003 | 0.999 | 0.010 | 0.306 | 0.368 | 0.4 |

| | | | | | | | | | | | | | | | | |
|---|---|---|---|---|---|---|---|---|---|---|---|---|---|---|---|---|
| Genic (±2 kb) | 2,266,807 | Gene struct. | Upstream (2K) | 831,210 | AL | 1.012 | 1.000 | 0.008 | 0.048 | 0.089 | 1.6 | 1.015 | 1.000 | 0.011 | 0.085 | 0.152 | 1.4 |
| Genic (±2 kb) | 2,266,807 | Gene struct. | Upstream (2K) | 831,210 | STR | 1.016 | 1.000 | 0.008 | 0.022 | 0.051 | 2.0 | 1.024 | 1.000 | 0.011 | 0.007 | 0.038 | 2.3 |
| Genic (±2 kb) | 2,266,807 | Gene struct. | 5' UTR | 87,319 | PHQ | 1.011 | 0.999 | 0.014 | 0.190 | 0.238 | 0.8 | 0.984 | 0.999 | 0.020 | 0.175 | 0.264 | -0.9 |
| Genic (±2 kb) | 2,266,807 | Gene struct. | 5' UTR | 87,319 | GSL | 1.021 | 1.000 | 0.013 | 0.057 | 0.103 | 1.6 | 1.012 | 1.000 | 0.020 | 0.244 | 0.331 | 0.6 |
| Genic (±2 kb) | 2,266,807 | Gene struct. | 5' UTR | 87,319 | AL | 1.004 | 1.000 | 0.014 | 0.397 | 0.417 | 0.3 | 1.001 | 0.999 | 0.019 | 0.471 | 0.471 | 0.1 |
| Genic (±2 kb) | 2,266,807 | Gene struct. | 5' UTR | 87,319 | STR | 1.011 | 1.000 | 0.013 | 0.204 | 0.252 | 0.8 | 1.004 | 0.999 | 0.019 | 0.401 | 0.428 | 0.2 |
| Genic (±2 kb) | 2,266,807 | Gene struct. | CDS | 241,038 | PHQ | 1.010 | 1.000 | 0.008 | 0.124 | 0.176 | 1.2 | 1.005 | 1.000 | 0.012 | 0.334 | 0.386 | 0.3 |
| Genic (±2 kb) | 2,266,807 | Gene struct. | CDS | 241,038 | GSL | 1.000 | 1.000 | 0.008 | 0.513 | 0.513 | 0.0 | 1.017 | 1.000 | 0.013 | 0.096 | 0.163 | 1.3 |
| Genic (±2 kb) | 2,266,807 | Gene struct. | CDS | 241,038 | AL | 1.007 | 1.000 | 0.009 | 0.223 | 0.268 | 0.8 | 1.019 | 1.000 | 0.012 | 0.077 | 0.140 | 1.5 |
| Genic (±2 kb) | 2,266,807 | Gene struct. | CDS | 241,038 | STR | 1.013 | 1.000 | 0.009 | 0.073 | 0.118 | 1.4 | 1.007 | 1.000 | 0.012 | 0.286 | 0.349 | 0.6 |
| Genic (±2 kb) | 2,266,807 | Gene struct. | Intron | 432,870 | PHQ | 0.994 | 1.000 | 0.011 | 0.303 | 0.346 | -0.5 | 0.993 | 0.999 | 0.015 | 0.379 | 0.410 | -0.5 |
| Genic (±2 kb) | 2,266,807 | Gene struct. | Intron | 432,870 | GSL | 0.981 | 1.000 | 0.010 | 0.021 | 0.049 | -1.8 | 0.989 | 1.000 | 0.016 | 0.238 | 0.327 | -0.7 |
| Genic (±2 kb) | 2,266,807 | Gene struct. | Intron | 432,870 | AL | 0.980 | 0.999 | 0.012 | 0.056 | 0.103 | -1.6 | 0.955 | 0.998 | 0.016 | 0.002 | 0.018 | -2.6 |

| | | | | | | | | | | | | | | | | | |
|---|---|---|---|---|---|---|---|---|---|---|---|---|---|---|---|---|---|
| Genic (±2 kb) | 2,266,807 | Gene struct. | Intron | 432,870 | STR | 0.973 | 1.000 | 0.011 | 0.005 | 0.021 | -2.4 | 0.964 | 1.000 | 0.015 | 0.003 | 0.024 | -2.4 |
| Genic (±2 kb) | 2,266,807 | Gene struct. | 3' UTR | 153,807 | PHQ | 0.997 | 0.999 | 0.010 | 0.408 | 0.417 | -0.2 | 1.029 | 0.999 | 0.015 | 0.039 | 0.084 | 1.9 |
| Genic (±2 kb) | 2,266,807 | Gene struct. | 3' UTR | 153,807 | GSL | 1.005 | 1.000 | 0.010 | 0.296 | 0.345 | 0.4 | 1.002 | 1.000 | 0.016 | 0.455 | 0.466 | 0.1 |
| Genic (±2 kb) | 2,266,807 | Gene struct. | 3' UTR | 153,807 | AL | 1.003 | 1.000 | 0.011 | 0.403 | 0.417 | 0.2 | 1.029 | 1.001 | 0.016 | 0.029 | 0.072 | 1.9 |
| Genic (±2 kb) | 2,266,807 | Gene struct. | 3' UTR | 153,807 | STR | 0.995 | 1.000 | 0.011 | 0.361 | 0.393 | -0.4 | 1.010 | 0.999 | 0.015 | 0.246 | 0.331 | 0.7 |
| Genic (±2 kb) | 2,266,807 | Gene struct. | Downstream (2K) | 520,563 | PHQ | 0.996 | 1.000 | 0.008 | 0.308 | 0.346 | -0.5 | 1.003 | 0.999 | 0.012 | 0.377 | 0.410 | 0.3 |
| Genic (±2 kb) | 2,266,807 | Gene struct. | Downstream (2K) | 520,563 | GSL | 1.003 | 1.000 | 0.008 | 0.313 | 0.349 | 0.5 | 0.991 | 1.000 | 0.011 | 0.204 | 0.295 | -0.8 |
| Genic (±2 kb) | 2,266,807 | Gene struct. | Downstream (2K) | 520,563 | AL | 0.995 | 1.001 | 0.009 | 0.242 | 0.289 | -0.7 | 0.998 | 1.001 | 0.012 | 0.415 | 0.434 | -0.2 |
| Genic (±2 kb) | 2,266,807 | Gene struct. | Downstream (2K) | 520,563 | STR | 0.995 | 1.000 | 0.009 | 0.297 | 0.345 | -0.6 | 0.994 | 0.999 | 0.012 | 0.335 | 0.386 | -0.5 |

**Table S3. Gene-level genomic contexts and enrichment statistics.**

Gene-level enrichment was evaluated using gene-level envGWAS results. Site-level association signals were first aggregated to genes using MAGMA, and genes were then ranked by association *P* value for each environmental variable.

**Category** and **Context** define the gene-level genomic context being evaluated, and **N in context** indicates the number of genes assigned to that context. **Envir.** indicates the environmental variable tested.

Enrichment was quantified as the Jaccard index between the context-defined gene set and the top 5% or top 1% most strongly associated genes. **Top5 obs.** and **Top1 obs.** report the observed Jaccard indices for the top 5% and top 1% gene sets, respectively. **Top5 null med.** / **Top1 null med.** and **Top5 null SD** / **Top1 null SD** summarize Jaccard indices across 1,000 equal-size random gene sets. **Top5 emp. P** / **Top1 emp. P** report empirical *P* values, and **Top5 BH FDR** / **Top1 BH FDR** report Benjamini–Hochberg adjusted *P* values. **Top5 Z** and **Top1 Z** are standardized enrichment scores relative to the null distribution; positive values indicate enrichment and negative values indicate depletion of GEA signals in the corresponding gene-level context.

Abbreviations: **PHQ**, precipitation of the warmest quarter; **GSL**, growing season length; **AL**, extractable aluminum; **STR**, *Striga* HS score.

| Category | Context | N in context | Envir. | Top5 obs. | Top5 null med. | Top5 null SD | Top5 emp *P* | Top5 BD FDR | Top5 *Z* | Top1 obs. | Top1 null med. | Top1 null SD | Top1 emp *P* | Top1 BD FDR | Top1 *Z* |
|---|---|---|---|---|---|---|---|---|---|---|---|---|---|---|---|
| Expression | No detected | 1,232 | PHQ | 0.015 | 0.023 | 0.003 | 0.005 | 0.078 | -2.5 | 0.004 | 0.008 | 0.002 | 0.034 | 0.442 | -1.8 |
| Expression | No detected | 1,232 | GSL | 0.015 | 0.022 | 0.003 | 0.004 | 0.078 | -2.4 | 0.005 | 0.008 | 0.002 | 0.149 | 0.459 | -1.2 |
| Expression | No detected | 1,232 | AL | 0.018 | 0.023 | 0.003 | 0.065 | 0.267 | -1.5 | 0.005 | 0.008 | 0.003 | 0.146 | 0.459 | -1.2 |
| Expression | No detected | 1,232 | STR | 0.024 | 0.023 | 0.003 | 0.419 | 0.505 | 0.2 | 0.008 | 0.008 | 0.002 | 0.509 | 0.551 | -0.2 |
| Expression | High var. expr. | 5,195 | PHQ | 0.043 | 0.040 | 0.002 | 0.110 | 0.312 | 1.3 | 0.012 | 0.010 | 0.001 | 0.100 | 0.459 | 1.4 |
| Expression | High var. expr. | 5,195 | GSL | 0.048 | 0.040 | 0.002 | 0.004 | 0.078 | 2.9 | 0.011 | 0.010 | 0.001 | 0.283 | 0.499 | 0.7 |
| Expression | High var. expr. | 5,195 | AL | 0.040 | 0.040 | 0.002 | 0.439 | 0.505 | -0.2 | 0.011 | 0.010 | 0.001 | 0.163 | 0.471 | 1.1 |

| | | | | | | | | | | | | | | | |
|---|---|---|---|---|---|---|---|---|---|---|---|---|---|---|---|
| Expression | High var. expr. | 5,195 | STR | 0.042 | 0.041 | 0.003 | 0.339 | 0.500 | 0.4 | 0.010 | 0.010 | 0.001 | 0.393 | 0.504 | 0.3 |
| Expression | Stably | 1,350 | PHQ | 0.024 | 0.024 | 0.003 | 0.458 | 0.505 | 0.1 | 0.008 | 0.008 | 0.002 | 0.437 | 0.504 | -0.3 |
| Expression | Stably | 1,350 | GSL | 0.021 | 0.023 | 0.003 | 0.229 | 0.411 | -0.8 | 0.005 | 0.008 | 0.002 | 0.095 | 0.459 | -1.3 |
| Expression | Stably | 1,350 | AL | 0.026 | 0.023 | 0.003 | 0.190 | 0.399 | 0.9 | 0.009 | 0.008 | 0.002 | 0.393 | 0.504 | 0.3 |
| Expression | Stably | 1,350 | STR | 0.027 | 0.024 | 0.003 | 0.170 | 0.399 | 1.0 | 0.007 | 0.008 | 0.002 | 0.402 | 0.504 | -0.4 |
| Recombination | High | 7,438 | PHQ | 0.052 | 0.043 | 0.006 | 0.102 | 0.312 | 1.3 | 0.012 | 0.010 | 0.002 | 0.251 | 0.489 | 0.7 |
| Recombination | High | 7,438 | GSL | 0.047 | 0.043 | 0.006 | 0.260 | 0.423 | 0.6 | 0.010 | 0.010 | 0.002 | 0.441 | 0.504 | 0.1 |
| Recombination | High | 7,438 | AL | 0.041 | 0.043 | 0.006 | 0.365 | 0.500 | -0.3 | 0.007 | 0.009 | 0.002 | 0.141 | 0.459 | -1.1 |
| Recombination | High | 7,438 | STR | 0.044 | 0.044 | 0.006 | 0.497 | 0.505 | 0.0 | 0.010 | 0.010 | 0.002 | 0.395 | 0.504 | 0.3 |
| Recombination | Med. | 19,992 | PHQ | 0.050 | 0.049 | 0.003 | 0.477 | 0.505 | 0.2 | 0.011 | 0.010 | 0.001 | 0.191 | 0.489 | 0.9 |
| Recombination | Med. | 19,992 | GSL | 0.052 | 0.050 | 0.003 | 0.177 | 0.399 | 1.0 | 0.011 | 0.010 | 0.001 | 0.273 | 0.499 | 0.7 |
| Recombination | Med. | 19,992 | AL | 0.054 | 0.049 | 0.003 | 0.057 | 0.267 | 1.6 | 0.012 | 0.010 | 0.001 | 0.026 | 0.442 | 2.0 |
| Recombination | Med. | 19,992 | STR | 0.053 | 0.050 | 0.003 | 0.110 | 0.312 | 1.2 | 0.011 | 0.010 | 0.001 | 0.239 | 0.489 | 0.7 |
| Recombination | Low | 4,247 | PHQ | 0.026 | 0.037 | 0.009 | 0.052 | 0.267 | -1.5 | 0.003 | 0.009 | 0.004 | 0.007 | 0.364 | -1.8 |
| Recombination | Low | 4,247 | GSL | 0.024 | 0.038 | 0.007 | 0.032 | 0.267 | -2.0 | 0.007 | 0.009 | 0.003 | 0.209 | 0.489 | -1.0 |
| Recombination | Low | 4,247 | AL | 0.025 | 0.038 | 0.007 | 0.038 | 0.267 | -1.9 | 0.006 | 0.009 | 0.003 | 0.106 | 0.459 | -1.2 |
| Recombination | Low | 4,247 | STR | 0.027 | 0.038 | 0.008 | 0.056 | 0.267 | -1.4 | 0.006 | 0.009 | 0.003 | 0.105 | 0.459 | -1.3 |
| Flowering | Flowering time genes | 312 | PHQ | 0.010 | 0.008 | 0.002 | 0.247 | 0.414 | 0.8 | 0.009 | 0.005 | 0.003 | 0.119 | 0.459 | 1.5 |
| Flowering | Flowering time genes | 312 | GSL | 0.014 | 0.008 | 0.002 | 0.009 | 0.094 | 2.6 | 0.006 | 0.005 | 0.003 | 0.430 | 0.504 | 0.4 |

| | | | | | | | | | | | | | | | |
|---|---|---|---|---|---|---|---|---|---|---|---|---|---|---|---|
| Flowering | Flowering time genes | 312 | AL | 0.009 | 0.008 | 0.002 | 0.312 | 0.492 | 0.6 | 0.005 | 0.005 | 0.003 | 0.631 | 0.631 | -0.1 |
| Flowering | Flowering time genes | 312 | STR | 0.005 | 0.008 | 0.002 | 0.072 | 0.267 | -1.5 | 0.005 | 0.005 | 0.003 | 0.628 | 0.631 | -0.1 |
| GCNs | Hub genes | 7,130 | PHQ | 0.041 | 0.043 | 0.002 | 0.124 | 0.322 | -1.2 | 0.009 | 0.010 | 0.001 | 0.231 | 0.489 | -0.8 |
| GCNs | Hub genes | 7,130 | GSL | 0.043 | 0.043 | 0.002 | 0.463 | 0.505 | 0.1 | 0.009 | 0.010 | 0.001 | 0.218 | 0.489 | -0.9 |
| GCNs | Hub genes | 7,130 | AL | 0.041 | 0.043 | 0.002 | 0.236 | 0.411 | -0.7 | 0.011 | 0.010 | 0.001 | 0.108 | 0.459 | 1.3 |
| GCNs | Hub genes | 7,130 | STR | 0.046 | 0.044 | 0.002 | 0.192 | 0.399 | 0.9 | 0.010 | 0.010 | 0.001 | 0.482 | 0.533 | 0.1 |
| GCNs | Non-hub module genes | 20,422 | PHQ | 0.050 | 0.049 | 0.001 | 0.138 | 0.342 | 1.2 | 0.011 | 0.010 | 0.000 | 0.123 | 0.459 | 1.2 |
| GCNs | Non-hub module genes | 20,422 | GSL | 0.049 | 0.050 | 0.001 | 0.401 | 0.505 | -0.3 | 0.011 | 0.010 | 0.000 | 0.242 | 0.489 | 0.7 |
| GCNs | Non-hub module genes | 20,422 | AL | 0.050 | 0.049 | 0.001 | 0.357 | 0.500 | 0.4 | 0.010 | 0.010 | 0.000 | 0.388 | 0.504 | -0.3 |
| GCNs | Non-hub module genes | 20,422 | STR | 0.051 | 0.050 | 0.001 | 0.375 | 0.500 | 0.3 | 0.011 | 0.010 | 0.000 | 0.125 | 0.459 | 1.2 |
| GCNs | Non-module genes | 4,267 | PHQ | 0.038 | 0.038 | 0.002 | 0.467 | 0.505 | -0.2 | 0.009 | 0.010 | 0.001 | 0.288 | 0.499 | -0.7 |
| GCNs | Non-module genes | 4,267 | GSL | 0.039 | 0.038 | 0.003 | 0.402 | 0.505 | 0.2 | 0.010 | 0.010 | 0.001 | 0.524 | 0.556 | 0.0 |
| GCNs | Non-module genes | 4,267 | AL | 0.039 | 0.038 | 0.003 | 0.363 | 0.500 | 0.4 | 0.008 | 0.009 | 0.001 | 0.150 | 0.459 | -1.1 |
| GCNs | Non-module genes | 4,267 | STR | 0.035 | 0.039 | 0.003 | 0.064 | 0.267 | -1.6 | 0.007 | 0.009 | 0.001 | 0.033 | 0.442 | -1.8 |
| Gene Family | Single-copy | 16,161 | PHQ | 0.048 | 0.048 | 0.002 | 0.490 | 0.505 | 0.0 | 0.010 | 0.010 | 0.001 | 0.406 | 0.504 | -0.3 |
| Gene Family | Single-copy | 16,161 | GSL | 0.048 | 0.049 | 0.002 | 0.375 | 0.500 | -0.3 | 0.011 | 0.010 | 0.001 | 0.201 | 0.489 | 0.9 |
| Gene Family | Single-copy | 16,161 | AL | 0.049 | 0.048 | 0.002 | 0.237 | 0.411 | 0.8 | 0.010 | 0.010 | 0.001 | 0.382 | 0.504 | 0.3 |
| Gene Family | Single-copy | 16,161 | STR | 0.050 | 0.049 | 0.002 | 0.216 | 0.411 | 0.8 | 0.010 | 0.010 | 0.001 | 0.254 | 0.489 | 0.8 |
| Gene Family | Multi-copy | 12,421 | PHQ | 0.047 | 0.047 | 0.002 | 0.505 | 0.505 | 0.0 | 0.011 | 0.010 | 0.001 | 0.397 | 0.504 | 0.3 |

| | | | | | | | | | | | | | | | |
|---|---|---|---|---|---|---|---|---|---|---|---|---|---|---|---|
| Gene Family | Multi-copy | 12,421 | GSL | 0.050 | 0.047 | 0.002 | 0.072 | 0.267 | 1.6 | 0.010 | 0.010 | 0.001 | 0.395 | 0.504 | -0.3 |
| Gene Family | Multi-copy | 12,421 | AL | 0.047 | 0.047 | 0.002 | 0.486 | 0.505 | -0.1 | 0.010 | 0.010 | 0.001 | 0.446 | 0.504 | -0.2 |
| Gene Family | Multi-copy | 12,421 | STR | 0.045 | 0.048 | 0.002 | 0.114 | 0.312 | -1.2 | 0.010 | 0.010 | 0.001 | 0.303 | 0.504 | -0.6 |
| Gene Family | Orphan | 3,237 | PHQ | 0.035 | 0.035 | 0.003 | 0.460 | 0.505 | 0.1 | 0.010 | 0.010 | 0.002 | 0.538 | 0.560 | 0.0 |
| Gene Family | Orphan | 3,237 | GSL | 0.028 | 0.035 | 0.003 | 0.006 | 0.078 | -2.5 | 0.008 | 0.009 | 0.002 | 0.150 | 0.459 | -1.1 |
| Gene Family | Orphan | 3,237 | AL | 0.031 | 0.035 | 0.003 | 0.083 | 0.288 | -1.4 | 0.009 | 0.009 | 0.002 | 0.410 | 0.504 | -0.3 |
| Gene Family | Orphan | 3,237 | STR | 0.038 | 0.036 | 0.003 | 0.235 | 0.411 | 0.7 | 0.009 | 0.009 | 0.002 | 0.401 | 0.504 | -0.4 |

**Table S4. Variance explained by genomic contexts.**

Variance partitioning was used to estimate the proportion of environmental variation explained by sites within focal genomic contexts.

**Context** indicates the genomic context used to construct the foreground GRM, and **Envir.** indicates the environmental variable used as the response. **Var. expl.** reports the foreground context-specific estimate, and **Genome var. expl.** reports the corresponding whole-genome estimate. Matched control GRMs were generated from random site sets matched for site number, MAF, recombination rate, and $\pi$. **Perm. ctrl. med.** and **Perm. ctrl. SD** report the median and standard deviation of variance explained across matched control GRMs. **Emp. *P*** is the empirical *P* value from 1,000 matched-site permutations, and **BH FDR** is the Benjamini–Hochberg adjusted *P* value. **FG *Z*** is the standardized deviation of the foreground estimate from the matched-control distribution; positive values indicate greater variance explained than expected for matched random site sets.

Abbreviations: **PHQ**, precipitation of the warmest quarter; **GSL**, growing season length; **AL**, extractable aluminum; **STR**, *Striga* HS score.

| Context | Envir. | Var. expl. | Genome var. expl. | Perm. ctrl. med. | Perm. ctrl. SD | Emp. *P* | BH FDR | FG *Z* |
|---|---|---|---|---|---|---|---|---|
| ACRs | PHQ | 0.615 | 0.549 | 0.609 | 0.007 | 0.190 | 0.207 | 0.9 |
| ACRs | GSL | 0.820 | 0.817 | 0.825 | 0.003 | 0.057 | 0.068 | -1.6 |
| ACRs | AL | 0.806 | 0.746 | 0.797 | 0.004 | 0.011 | 0.015 | 2.2 |
| ACRs | STR | 0.693 | 0.665 | 0.678 | 0.004 | 0.001 | 0.002 | 3.6 |
| UMRs | PHQ | 0.621 | 0.549 | 0.606 | 0.001 | 0.001 | 0.002 | 10.2 |
| UMRs | GSL | 0.831 | 0.817 | 0.824 | 0.001 | 0.001 | 0.002 | 9.9 |
| UMRs | AL | 0.803 | 0.746 | 0.795 | 0.001 | 0.001 | 0.002 | 9.4 |
| UMRs | STR | 0.691 | 0.665 | 0.671 | 0.001 | 0.001 | 0.002 | 18.3 |
| Proximal TEs | PHQ | 0.641 | 0.549 | 0.567 | 0.003 | 0.001 | 0.002 | 18.8 |
| Proximal TEs | GSL | 0.821 | 0.817 | 0.822 | 0.002 | 0.295 | 0.295 | -0.6 |
| Proximal TEs | AL | 0.789 | 0.746 | 0.766 | 0.002 | 0.001 | 0.002 | 9.5 |
| Proximal TEs | STR | 0.697 | 0.665 | 0.668 | 0.002 | 0.001 | 0.002 | 12.5 |

**Table S5. Genomic prediction of drought response.**

Genomic prediction was used to test whether growing season length–associated sites within focal genomic contexts could predict grain-yield response to drought in the Brazil drought trials.

**Context** indicates the genomic context used to construct the foreground GRM. **Pred.** reports prediction accuracy for drought response based on the foreground GRM, measured as the Pearson correlation between predicted growing season length genetic scores and grain-yield plasticity. **Genome pred.** reports the corresponding prediction accuracy based on the whole-genome GRM.

Matched control GRMs were generated from random site sets matched for site number, MAF, recombination rate, and $\pi$. **Perm. ctrl. med.** and **Perm. ctrl. SD** reports the median and standard deviation of prediction accuracy across matched control GRMs. **Emp.** ***P*** is the empirical *P* value from 1,000 matched-site permutations. **FG** ***Z*** is the standardized deviation of the foreground prediction accuracy from the matched control distribution; positive values indicate greater predictive accuracy than expected for matched random site sets.

| Context | Pred. | Genome pred. | Perm. ctrl. med. | Perm. ctrl. SD | Emp. *P* | FG *Z* |
|---|---|---|---|---|---|---|
| ACRs | 0.292 | 0.280 | 0.271 | 0.007 | 0.003 | 3.0 |
| UMRs | 0.276 | 0.280 | 0.268 | 0.001 | 0.001 | 5.9 |
| Proximal TEs | 0.275 | 0.280 | 0.284 | 0.003 | 0.004 | -2.8 |

**Dataset S1. Accession metadata**
**Dataset S2. *Striga* occurrence records**
**Dataset S3. Environmental predictors for *Striga* SDM**
**Dataset S4. Flowering-time genes and *cis*-regulatory adaptation genes**
**Dataset S5. Phenotypic datasets and trait values**
**Dataset S6. Metadata for RNA-seq samples used in Gene coexpression networks construction**
**Dataset S7. Gene coexpression network modules**
**Dataset S8. Genes with significant *cis*-eQTLs**
